# From Stress to Survival: Trophoblast-Derived Extracellular Vesicle Proteome Captures Aspirin-Driven Cellular Reprogramming in a Preeclampsia Model

**DOI:** 10.64898/2026.03.08.710202

**Authors:** Vineet Mahajan, Awanit Kumar, Jeena Jacob, Maged Costantine, Lauren S. Richardson, Rheanna Urrabaz-Garza, Emmanuel Amabebe, Ourlad A. Tantengco, Ananth Kumar Kammala, Ramkumar Menon

## Abstract

**Background:** Low-dose aspirin (LDA) reduces preeclampsia (PE) risk by up to 40%, yet its molecular effects on chorion trophoblast cells (CTCs) a fetal membrane lineage at the feto-maternal interface remain obscure. CTCs form a structural and immunoregulatory barrier whose dysfunction drives inflammation-associated membrane pathology in PE. Extracellular vesicles (EVs) released by CTCs may encode cellular stress and adaptation states, offering a molecular window into aspirin’s timing-dependent effects on PE risk modification.

**Methods:** Human CTCs were challenged with cigarette smoke extract (CSE) to model oxidative stress-driven PE pathology. Two paradigms were tested: (1) prophylactic aspirin (4 and 40 µg/ml) before and/or flanking CSE, and (2) therapeutic aspirin after CSE challenge. EVs were isolated via ultracentrifugation and size exclusion chromatography, characterized by nanoparticle tracking and immunoblotting, and profiled by quantitative mass spectrometry. Network pathway analysis and machine-learning biomarker selection defined EV-encoded molecular states.

**Results:** CTC-derived EVs from CSE-exposed cells carried a PE-like proteomic signature marked by suppressed VEGF/ECM remodeling, activated TNF-p53 apoptotic signaling, and heightened inflammation. Prophylactic low-dose aspirin shifted EV cargo toward preserved angiogenic capacity (VEGFA, COL1A1, MMP14) with attenuated apoptotic and NF-κB signatures. High-dose aspirin produced broad transcriptional suppression without vascular benefit in EVs. Therapeutic aspirin partially rescued injury-associated EV cargo but failed to restore angiogenic signatures. Machine-learning analysis of EV proteomes identified a prophylactic biomarker panel anchored by HSPA8, SERPINF2, COL4A1, and PLOD1, linked to angiogenic recovery and redox balance.

**Conclusions:** CTC-derived EV proteomic signatures capture dose-and timing-dependent aspirin effects, positioning the chorion as a pharmacological “secondary responder” favoring cellular resilience over classical anti-inflammatory suppression. EV-based molecular profiling might offer a framework for stratifying aspirin responders from non-responders toward personalized PE prevention.

## 1. Introduction

Preeclampsia (PE), together with other placental pathologies of pregnancy including fetal growth restriction (FGR), is a major contributor to maternal and neonatal morbidity worldwide.^1,2,3,4^

The placenta and specifically the trophoblast layer is a primary target of oxidative stress, which impair trophoblast invasion, disrupt vascular remodeling, and dysregulate immune tolerance at the feto-maternal interface.^5,6^

Low-dose aspirin (LDA) prophylaxis remains the gold standard for high-risk pregnancies, reducing the incidence of preeclampsia by approximately 40% when initiated before 16 weeks of gestation. ^7,8,9^ However, this leaves a substantial proportion of patients as’non-responders,’ a clinical reality that current placenta-centric models fail to explain.^10,11,12^ The exact targets of aspirin’s action, as well as its timing-dependent effects and mechanisms of action at the feto-maternal interface, remain unclear despite its widespread medical use.

The feto-maternal interface comprises two distinct compartments the placenta/decidua, and the fetal membrane/ decidua each contributing distinct functions to maintain pregnancy. Within this interface, chorion trophoblast cells (CTCs) form a key mechanical and immunological barrier, regulating biochemical exchange, ECM remodeling, and choriodecidual angiogenic signaling during pregnancy.^13,14^ Dysregulation of these functions, primarily mediated by PE associated inflammation and oxidative stress, has been implicated in adverse pregnancy outcomes resulting in preterm birth that often coexist with placental dysfunction.^15,16^ The interventions for adverse pregnancy outcomes are often developed based on the mechanisms of action of compounds studied in placental trophoblast cells. As CTCs are the same lineage of placental alkaline phosphatase (PLAP) ^17^ expressing trophoblast cells, with similar functions, it is crucial to know how CTCs contribute to better the outcome after a treatment.^18^ Understanding how CTCs respond to aspirin treatment is therefore essential to define whether aspirin confers systemic protection across all feto-maternal interfaces (placenta/decidua and choriodecidua) or act primarily through placental pathways.

Although aspirin’s protective effects on placental endothelial and trophoblast signaling have been previously explored ^19,20,21^ its impact on fetal membrane trophoblasts, particularly CTCs, remains largely unexplored. Like placental villous trophoblasts, CTCs experience unique mechanical and oxidative stressors at the choriodecidual interface, where localized inflammation and extracellular vesicle (EV; exosomes of 30 – 200 nm) communication modulate tissue integrity and immune tolerance. In pregnancy, trophoblast-derived EVs are released into the maternal circulation from early gestation and play critical roles in immune modulation, angiogenesis, and spiral artery remodeling. ^22,23,24^ Alterations in circulating placental EV cargo have been reported in PE ^25,26^ and FGR ^27^ establishing EVs as both potential mediators of placental pathology and candidate liquid biopsy biomarkers. While the protective effects of aspirin on placental angiogenesis are well-documented, the pharmacological sensitivity of the CTC layer the critical immunoregulatory and structural barrier at the feto-maternal interface remains virtually uncharacterized. Importantly, CTCs express PLAP, enabling identification of CTC-derived EVs in maternal circulation through PLAP-based capture strategies.^28^ The choriodecidual interface exposes CTCs to specific mechanical forces and oxidative stress because they exist in a distinct environment where local inflammation and EV communication systems affect tissue stability and immune acceptance. The timing of aspirin exposure in relation to inflammatory or oxidative stress remains a critical question for maternal-fetal pharmacology. The protective effects of aspirin are expected to emerge from its administration that creates an anti-inflammatory state through COX-dependent eicosanoid signaling and NF-κB suppression, following pre-inflammatory stressor administration.^29,30^ The protective effect of preconditioning helps maintain chorionic cell survival and supports the production of angiogenic factors and provide membrane stability. In contrast, therapeutic aspirin treatment, where aspirin is introduced after an insult, may act through secondary mechanisms limiting damage propagation, restoring redox balance, or promoting reparative signaling.^31, 32^

Differentiating these timing-dependent effects within CTC is critical because aspirin is often initiated at variable gestational stages in clinical practice, and its prophylactic versus therapeutic efficacy may diverge between placental and fetal membrane compartments. Given the central role of the chorion in maintaining fetal-maternal barrier integrity, deciphering aspirin’s direct and EV-mediated effects on CTC signaling, angiogenesis, apoptosis, and inflammation is essential. This mechanism would 1) Define whether aspirin’s benefits extend beyond the placenta to the fetal membranes 2) Clarify if prophylactic and therapeutic exposures elicit distinct molecular outcomes 3) Inform and identify gestational timing strategies to optimize aspirin use in reducing both preeclampsia and preterm birth.

This study is focused on investigating how low-dose aspirin, when administered either before (prophylactic/risk-exposure) or after (therapeutic) oxidative injury, alters the proteomic and functional landscape of CTC-derived EVs. We postulate that the chorion functions not merely as a passive bystander, but as a distinct’secondary responder’ tissue. Here, we have investigated whether EV cargo derived from CTCs captures distinct molecular states corresponding to oxidative injury, partial recovery, and stress-adaptive resilience. Because EVs integrate molecular signals across time and stress states, they offer a unique window into cellular adaptation that cannot be captured by single time-point cytokine or transcript measurements. By profiling EV cargo, we can resolve whether trophoblasts encode injury, partial recovery, or stress-tolerant adaptive programs in response to oxidative challenge. Thus, EVs serve not merely as biomarkers but as molecular records of stress history and adaptive capacity at the feto-maternal interface. Aspirin exposure, here, is employed not as a therapeutic endpoint but also as a pharmacologic perturbation to reveal how EV proteomic composition reflects the timing of stress exposure and adaptive signaling. By integrating EV proteomics with network-based and machine-learning analyses, we define EV-encoded molecular states that distinguish injury-dominant from stress-resilient trophoblast responses, providing a framework for EV-based biomarker discovery at the feto-maternal interface.

## 2. Materials and Methods

### 2.1 Cell source and culture conditions

CTCs were isolated in-house from fetal membrane tissues obtained from John Sealy Hospital at the University of Texas Medical Branch (UTMB), Galveston, TX, USA. Tissues were collected under IRB-approved exempt protocols for discarded and de-identified specimens from uncomplicated term Cesarean deliveries. Isolated CTCs were immortalized and maintained as previously described. Cells were cultured in a 1:1 mixture of DMEM and Ham’s F-12 medium (Corning, Ref# 10-092-CV) supplemented with 0.2% fetal bovine serum, 0.1 mM 2-mercaptoethanol, 0.5% penicillin-streptomycin, 0.3% BSA, 1% ITS-X supplement, 2 μM CHIR99021, 0.5 μM A83-01, 1 μM SB431542, 1.5 μg/mL L-ascorbic acid, 50 ng/mL EGF, 0.8 mM valproic acid, and 5 μM Y-27632. Cells were subcultured at low density (approximately 2 × 10D cells per T175 flask) to avoid confluence at the time of harvest.

### 2.2 Immunocytochemistry

CTC identity was confirmed by immunocytochemistry for cytokeratin-7 (CK7) and placental alkaline phosphatase (PLAP). After 24 h of subculture, cells were fixed with 4% paraformaldehyde, permeabilized with 0.5% Triton X-100, and blocked with 3% BSA in PBS. Cells were incubated overnight at 4 °C with anti-CK7 (ab9021, Abcam) and anti-PLAP (ab243731, Abcam) antibodies (1:200). Following PBS washes, secondary antibodies anti-mouse Alexa Fluor 488 (ab150073) and anti-rabbit Alexa Fluor 594 (ab150080) were applied (1:1000, 1 h, room temperature, dark). Nuclei were counterstained with DAPI and mounted using MoWiol 4-88. Images were acquired on a Keyence fluorescence microscope (Itasca, IL).

### 2.3 Cigarette smoke extract (CSE) preparation

Water-soluble CSE was prepared as previously reported and diluted 1:50 in culture medium immediately prior to use to induce oxidative stress. ^33, 34^

### 2.4 Therapeutic aspirin treatment

Based on reported plasma Cmax of low-dose aspirin in pregnancy, 4 μg/mL and 40 μg/mL doses were selected.^35^ Aspirin (Sigma-Aldrich, Cat#1044006) stock was prepared at 10 mg/mL in DMSO and diluted in medium to working concentrations. CTCs were first exposed to CSE for 24 h (day 1–2), followed by addition of aspirin (day 2–3) with or without continued CSE. Control conditions included media alone, equivalent DMSO, and aspirin alone. After 24 h of aspirin treatment, conditioned culture media (CCM) and cell lysates were collected. Cells were lysed in 1× RIPA buffer containing protease and phosphatase inhibitors and stored at −20 °C.

### 2.5 Prophylactic aspirin treatment

For prophylactic studies, CTCs were treated with aspirin (4 or 40 μg/mL) for 24 h after subculture. Cells were then challenged with CSE (1:50) in the presence or absence of the same aspirin dose for an additional 48 h. A continuous exposure arm (Asp→CSE→Asp) was included. Controls received equivalent DMSO. CCM and lysates were harvested as above.

### 2.6 Cytokine measurement and Inflammation Index

IL-6, IL-8, IL-10, and TNF concentrations in CCM were quantified using multiplex immunoassays according to the manufacturer’s instructions (Millipore-Milliplex Cytokine Chemokine Growth Factor Panel A-Catalogue no HCAYTA-60K). To evaluate net inflammatory balance, a composite Inflammation Index was calculated for each replicate as: z(IL-6) + z(IL-8) + z(TNF) − z(IL-10).IL-6/IL-10 and TNF/IL-10 ratios were used as secondary metrics. For groupwise analyses and figure generation, individual biological replicates were assigned to treatment categories based on experimental sequence rather than numeric dose labels. All samples annotated as CSE+Asp1, CSE+Asp2, and CSE+Asp3—representing CSE exposure followed by aspirin treatment at either 4 or 40 μg/mL—were pooled into a single Asp+CSE group, as these conditions reflect the same biological paradigm of aspirin administered after oxidative stress. Samples receiving aspirin before and after CSE challenge (Asp→CSE→Asp) were designated as the “Continuous” group. Controls included Ctrl (untreated) and CSE alone; wells treated with aspirin in the absence of CSE were classified as Asp alone. The composite Inflammation Index and IL-6/IL-10 ratios were calculated for each replicate and summarized as mean ± SEM within these predefined groups (n = 3 per group). This grouping scheme was applied uniformly to both therapeutic and prophylactic datasets to enable direct comparison of aspirin timing effects.

### 2.7 Cytotoxicity and viability

Aspirin cytotoxicity (0–100 μg/mL) was assessed using LDH release (Abcam ab197004). Ten microliters of media were mixed with 90 μL LDH reaction mix, incubated 10 min, and fluorescence measured at Ex/Em 535/587 nm. Viability was assessed using AlamarBlue (Invitrogen); resazurin conversion was measured at Ex/Em 560/590 nm after 3 h incubation.

### 2.8 NF-**κ**B and p38 MAPK signaling

Protein levels of total and phospho-NF-κB and p38 MAPK were measured using Simple Western JESS™ (Protein Simple). Lysates (0.2 mg/mL) were analyzed with antibodies: NF-κB (#4764S), phospho-NF-κB (S536), p38 MAPK (#9212), phospho-p38 MAPK (#4511) (Cell Signaling; 1:50). Detection used the anti-rabbit chemiluminescent module (Bio-Techne DM001) and Compass software v6.3.

### 2.9 Extracellular vesicle isolation and characterization

EVs were isolated from CCM by differential centrifugation, 100 kDa ultrafiltration, 0.2 μm filtration, and ultracentrifugation at 100,000×g for 2 h at 4 °C, followed by ExoSpin purification. Particle size and concentration were determined by ZetaView PMX 110 (Particle Metrix). EV protein markers (PLAP, FLOT-1, CD9) were assessed using Simple Western JESS as above. Per MISEV2023, at least one transmembrane protein (CD9) and one cytosolic-recovered protein (FLOT-1) were assessed. ^36^ PLAP was included as a tissue-of-origin marker to confirm trophoblast derivation of the isolated EVs. We have submitted all relevant data of our experiments to the EV-TRACK knowledgebase^36^ (EV-TRACK ID: EV260015) and is currently being processed.

### 2.10 NanoLC-MS/MS proteomics

EV peptides were analyzed on an UltiMate 3000 RSLCnano coupled to an Orbitrap Eclipse using DIA with staggered 8-Da windows (400–900 m/z). Data were processed in FragPipe v22.0 with MSFragger/DIA-NN against UniProt Homo sapiens (2025-02-05). FDR was set to 2%; differential expression used limma with Benjamini–Hochberg correction (adjusted p < 0.05; |log2FC| ≥ 1).

### 2.11 Proteomics pathway analysis using Ingenuity Pathway Analysis (IPA)

Differentially expressed proteins identified from DIA proteomics were further interrogated using Ingenuity Pathway Analysis (IPA, Qiagen, Redwood City, CA) to determine enriched biological functions, canonical pathways, and upstream regulators. ^37^ Protein identifiers with corresponding log_₂_ fold changes and adjusted p-values were uploaded into IPA, and the Core Analysis module was performed using the Ingenuity Knowledge Base (human reference). Activation or inhibition of pathways and regulators was inferred using the z-score algorithm, with significance determined by right-tailed Fisher’s exact test (p < 0.05). Networks related to inflammatory signaling, oxidative stress, autophagy, and extracellular vesicle biogenesis were prioritized based on relevance to CTC biology and aspirin response. Predicted upstream regulators and mechanistic networks were integrated with cytokine and Inflammation Index data to generate hypothesis-driven models.

### 2.12 Omics Playground for integrative visualization and biomarker discovery

Raw protein abundance matrices generated from FragPipe/DIA-NN were imported into Omics playground (BigOmics, Switzerland) for complementary exploratory analysis ^38, 39^. Data were log_₂_-transformed, normalized and median-centered prior to analysis. The platform was used to perform principal component analysis (PCA), hierarchical clustering, biomarker discovery and pathway enrichment based on Reactome and Gene Ontology databases. ML-based biomarker discovery was performed using Omics Playground (BigOmics Analytics, Switzerland) [21]. Ensemble analysis of LASSO, Elastic Net, Random Forest, and PLS-DA was applied to identify proteins most strongly discriminating against CSE vs Asp+CSE and Continuous groups. Model performance was assessed by k-fold cross-validation with AUC-ROC. Top candidate biomarkers were cross-validated with IPA-derived pathways to prioritize EV proteins potentially mediating aspirin-dependent modulation of CTC responses. ML candidates were filtered for Protein secretion/shedding potential, Known plasma detectability, Commercial immunoassay availability, FBS contamination risk, Trophoblast/placental expression evidence.

### 2.13 Statistical analysis

Experiments included three biological replicates (n = 3). Groupwise Inflammation Index and cytokine ratios were summarized as mean ± SEM. One-way ANOVA with appropriate post-hoc tests was performed using GraphPad Prism; p < 0.05 was considered significant.

## 3. Results

Oxidative stress-associated cellular changes and sterile inflammatory mimicking the reported pathologies of PE were recreated using CTCs after treating them with cigarette smoke extract (CSE), a well-established oxidative stress inducer. This study incorporated two distinct experimental paradigms: a prophylactic arm, in which aspirin is administered before and after the oxidative stress exposure, and a therapeutic arm, in which aspirin is administered only after the onset of oxidative stress–induced injury in CTCs. This study tested two prophylactic strategies and one treatment option in vitro. The prophylactic strategies were as follows: (1) Asp (4 [low dose] and 40 µg/ml [high dose]) followed by CSE treatment, (Asp4–CSE-Asp4, Asp40-CSE-Asp40) (2) Asp followed by CSE treatment and continuing treatment with Asp [CSE-Asp 4 (4 mg/ml) and CSE-Asp 40 (40 µg /ml] as outlined in Figure 1. The treatment condition was also tested, where exposure to CSE was followed by treatment (CSE-ASP; both at 4 and 40 mg/ml). The data detailed below are based on our observations in EVs released from CTCs after exposure or treatments with the intent of developing trophoblast EVs (PLAP+ve) as a biomarker of feto-placental well-being during pregnancy. For clarity and ease of discussion, the abbreviations in parentheses will be used in the subsequent sections.

**Figure 1.**
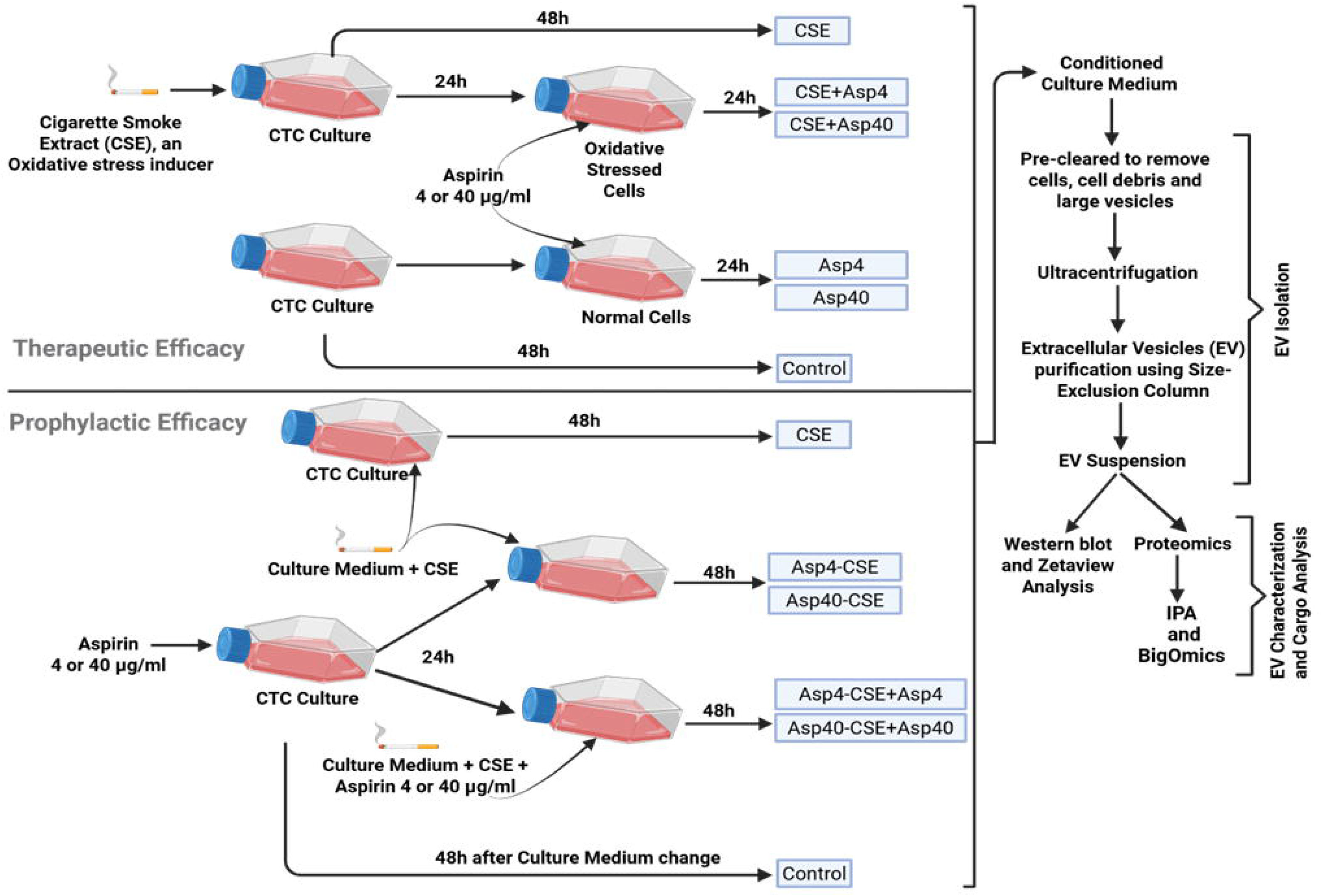
Experimental design illustrating therapeutic and prophylactic aspirin efficacy in oxidative stress induced chorion trophoblast cells and downstream EV)analysis. CTCs were cultured and exposed to cigarette smoke extract (CSE) to induce oxidative stress. Upper panel (Therapeutic Efficacy): CTCs were first subjected to CSE for 24 h, followed by treatment with aspirin (4 or 40 µg/mL; Asp4 or Asp40) for an additional 24 h. Parallel groups included CSE-only, aspirin-only (Asp4 or Asp40), and untreated control conditions. Lower panel (Prophylactic Efficacy): CTCs were co-treated with aspirin and CSE for 48 h (Asp4-CSE, Asp40-CSE),or pretreated with aspirin for 24 h prior to CSE exposure with continued aspirin (Asp4-CSE-Asp4, Asp40-CSE-Asp40) for 48 h. Conditioned culture media were collected 48 h after treatment or medium change, depending on group allocation. Right panel (EV workflow): Conditioned media were sequentially pre-cleared to remove cells, debris, and large vesicles, followed by ultracentrifugation and EV purification using size-exclusion chromatography. Purified EVs were characterised by ZetaView nanoparticle tracking analysis and Western blotting, and EV cargo was analysed using quantitative proteomics followed by Ingenuity Pathway Analysis (IPA) and Omics Playground machine learning. This experimental framework enables comparative assessment of aspirin’s therapeutic versus prophylactic effects on oxidative stress responses and EV-mediated signaling at the feto-maternal interface.

### 3.1 Cell characteristics

#### CTCs express Trophoblast markers and produce PLAP^+^ EVs across baseline and all treatment conditions

To validate the trophoblast phenotype of the CTCs, we first performed immunocytochemical and functional characterization assays prior to any CSE or aspirin treatment. CTCs showed distinct expression of trophoblast markers, with strong cytoplasmic staining for CK7 and membrane-associated expression of placental alkaline phosphatase (PLAP) (Fig. 2A). Cells formed cohesive epithelial monolayers with uniform nuclear morphology, confirming a stable trophoblast identity.

**Figure 2.**
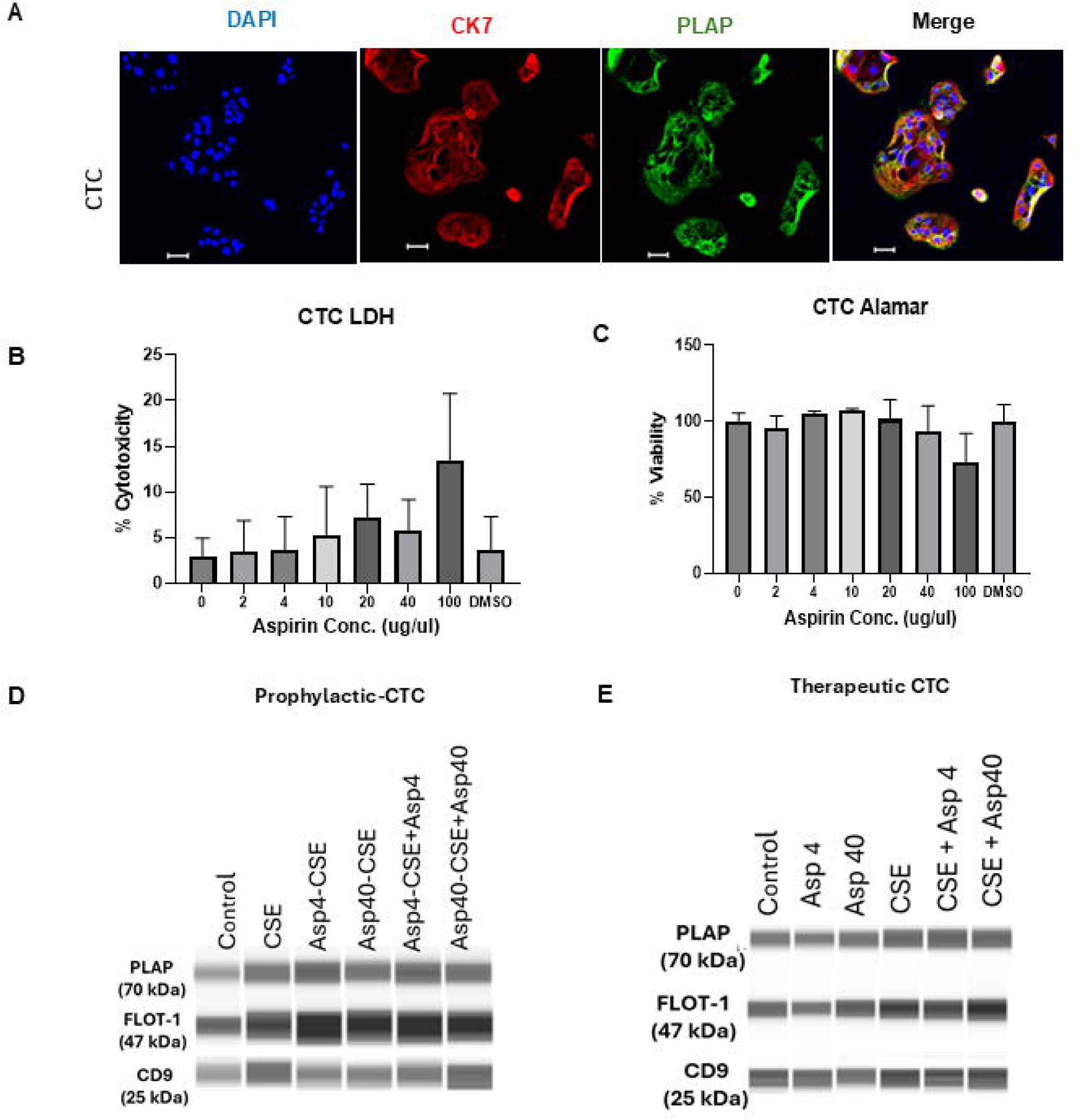
Cell characterisation and aspirin sensitivity. (A) Immunocytochemistry confirming co-expression of CK7 (red) and PLAP (green) in chorion trophoblast cells (CTCs), with DAPI nuclear counterstain (blue). Merged images confirm trophoblast identity. Scale bar: 50 µm. (B) LDH cytotoxicity assay across aspirin concentrations (0–100 µg/mL). Cytotoxicity remained <5% at 2–40 µg/mL, with modest increase (∼15%) at 100 µg/mL. DMSO vehicle control showed no cytotoxicity. (C) Alamar Blue viability assay. Viability was maintained at ∼100% across 2–40 µg/mL, with slight reduction at 100 µg/mL. (D) Western blotting of EVs isolated from prophylactic treatment conditions (Control, CSE, Asp4-CSE, Asp40-CSE, Asp4-CSE-Asp4, Asp40-CSE-Asp40), confirming expression of PLAP (70 kDa), FLOT-1 (47 kDa), and CD9 (25 kDa). (E) Western blotting of EVs from therapeutic conditions (Control, Asp4, Asp40, CSE, CSE+Asp4, CSE+Asp40), confirming PLAP, FLOT-1, and CD9 expression. PLAP detection in EVs from all conditions confirms trophoblast origin. Data represent mean ± SEM of n = 3 biological replicates.

Next, we assessed baseline cell viability and cytotoxicity across a range of aspirin concentrations to confirm dose tolerability. LDH assays showed low cytotoxicity across all doses tested, with values remaining near baseline up to 40 μg/μL and only mild elevation at 100 μg/μL (Fig. 2B). Alamar Blue assays showed cell viability (>90%) across all aspirin concentrations, indicating that CTCs tolerate low-to mid-range dosing without loss of metabolic activity (Fig. 2C).

Western blotting of EVs from all prophylactic conditions (Figure 2D, E) and all therapeutic conditions (Figure 2E) confirmed expression of EV markers FLOT-1 (47 kDa) and CD9 (25 kDa). Critically, PLAP (70 kDa) was detected in EVs from every treatment condition in both experimental arms, confirming that CTC-derived EVs retain trophoblast-origin identity regardless of CSE or aspirin exposure.

### 3.2 EV characterization under control and CSE exposure

EVs were isolated from conditioned media of all treatment groups by sequential ultracentrifugation and SEC purification, and characterised in accordance with MISEV2023 guidelines. EVs released by CTCs under baseline and CSE-exposed conditions were characterized using nanoparticle tracking analysis to define vesicle abundance and size distribution prior to downstream proteomic assessment. Across all conditions, EV preparations displayed size profiles consistent with small EVs (exosomes), with modal diameters ranging from 100–150 nm.

In the prophylactic dataset (summarized in Table1), EVs from untreated CTCs exhibited a mean concentration of 2.30×10¹D particles/mL and a mean diameter of 119.5 nm, representing the typical vesicle population produced by unstressed trophoblasts. EVs collected after CSE exposure showed a concentration of 4.50×10¹D particles/mL with a mean diameter of 105.2 nm, reflecting a shift toward a higher number of small EVs within the secreted population.

In the therapeutic dataset (summarized in Table 2), EV concentration remained within the same order of magnitude across conditions (ranging from 2.10×10¹D to 3.50×10¹D particles/mL), and mean EV diameters (approximately 135–142 nm) remained consistent with the definition of EVs or exosomes. These measurements indicate that CTCs maintain stable EV biophysical profiles during sustained exposure, with particle abundance varying within physiological ranges.

**Table 1:** Prophylactic_EV Characterization by ZetaView:

|  | Concentration<br>(Particles/ml) |  | Size<br>(nm) |  |
| --- | --- | --- | --- | --- |
|  | Mean | SD | Median | SD |
| **CTC Control** | 2.30E+10 | 2.0E+9 | 119.5 | 44.1 |
| **CTC_CSE** | 4.5E+10 | 6.0E+9 | 105.2 | 40.3 |
| **CTC_Asp4_CSE** | 5.8E+10 | 6.7E+9 | 107.8 | 46.3 |
| **CTC_Asp40_CSE** | 3.4E+10 | 4.0E+9 | 110.3 | 45.5 |
| **CTC_Asp4_CSE+Asp4** | 4.8E+10 | 1.2E+10 | 105.7 | 42 |
| **CTC_Asp40_CSE+Asp40** | 4.9E+10 | 5.5E+9 | 110.0 | 42.5 |

**Table 2:** Therapeutic EV Characterization by ZetaView.

|  | Concentration<br>(Particles/ml) |  | Size<br>(nm) |  |
| --- | --- | --- | --- | --- |
|  | Mean | SD | Median | SD |
| **CTC Control** | 2.60E+10 | 1.90E+09 | 141.7 | 61.9 |
| **CTC_Asp4** | 1.40E+10 | 1.2E+9 | 137.0 | 61.4 |
| **CTC_Asp40** | 2.60E+10 | 2.20E+09 | 135.8 | 58.0 |
| **CTC_CSE** | 2.40E+10 | 2.10E+09 | 142.4 | 64.4 |
| **CTC_CSE+Asp4** | 3.50E+10 | 3.60E+09 | 139.8 | 66.7 |
| **CTC_CSE+Asp40** | 2.10E+10 | 1.50E+09 | 138.0 | 54.1 |

### 3.3 CSE exposure establishes a preeclampsia-like baseline injury state and rewires the CTC-EV Proteome toward apoptosis, coagulation, and vascular suppression

To establish a baseline injury state for subsequent treatment comparisons, we first examined differential expression of proteins between untreated control and CSE-exposed CTC-derived EVs (Fig. 3A). Exposure to CSE drove a distinct shift in EV cargo composition. Among the most prominently upregulated proteins, Coagulation Factor X (F10) stood out as the single most significantly enriched hit, pointing toward a pro-coagulant EV phenotype under oxidative stress. This was accompanied by upregulation of Ribophorin I (RPN1), an endoplasmic reticulum stress marker, and CYP1A1, a well-characterized xenobiotic metabolism enzyme and classical CSE-responsive gene. Additional upregulated cargo included cytoskeletal stress marker Tubulin β1 (TUBB1), the metabolic signaling receptor INSR, and microtubule-associated protein MAP4 collectively suggesting broad cellular stress responses imprinted onto the EV proteome.

**Figure 3.**
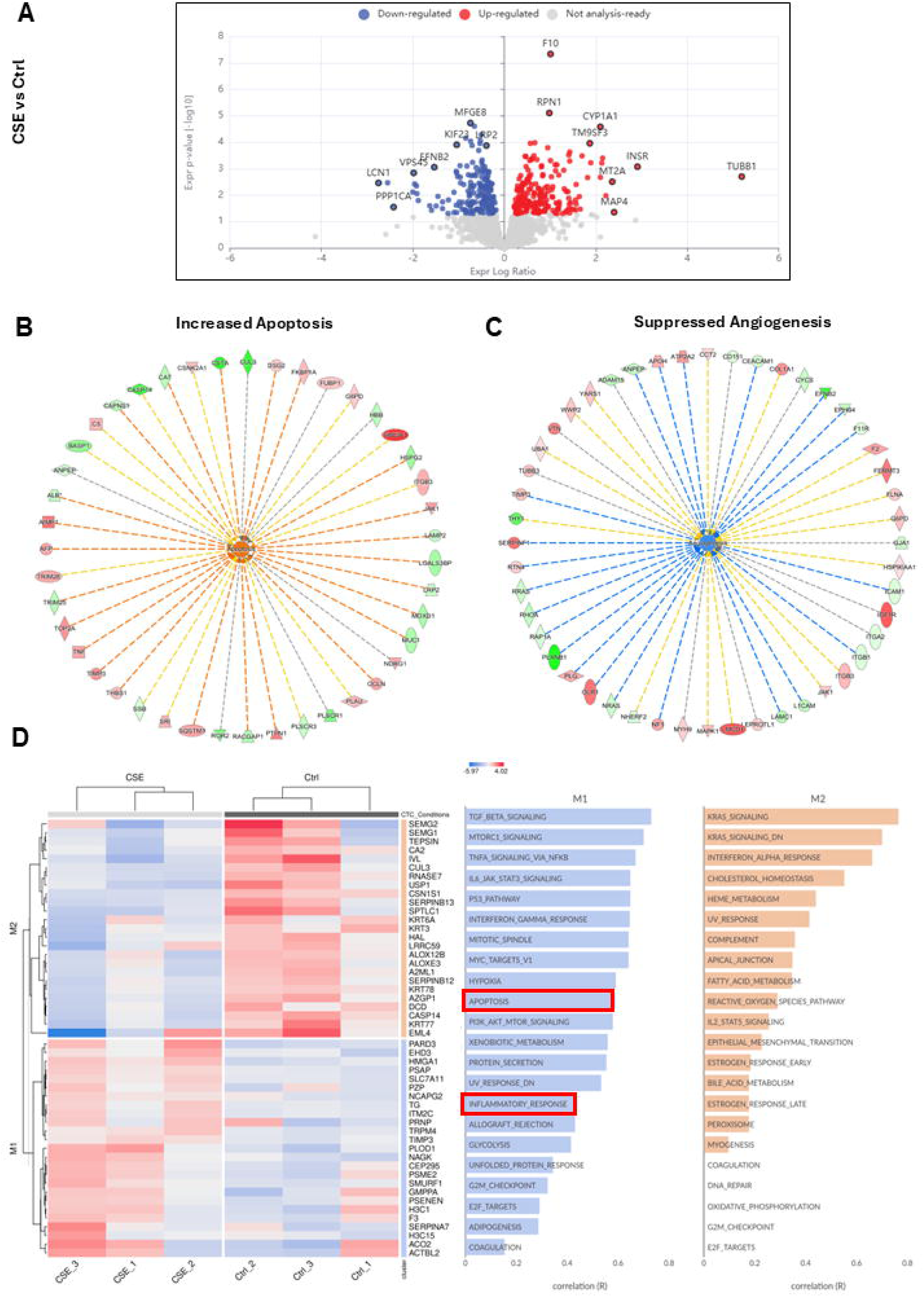
Oxidative stress rewires CTC-EV proteomic signatures toward cell death and vascular suppression. (A) Volcano plot showing differential protein expression in CSE versus control CTC-EVs. Red: upregulated; blue: downregulated; grey: not significant or not analysis-ready. F10 (Coagulation Factor X) was the most significantly upregulated protein; MFG-E8 (Lactadherin) was among the most significantly downregulated pro-angiogenic proteins. (B) IPA functional network predicting increased apoptosis in CSE-derived EVs. (C) IPA functional network predicting reduced angiogenesis in CSE-derived EVs. (D) Left: Hierarchical clustering of differentially expressed EV proteins between CSE and control conditions, showing clear sample segregation. Right: Pathway enrichment analysis. Module 1 (M1, blue bars): pathways positively correlated with the CSE-altered proteome, including TGF-β, mTORC1, TNFα/NF-κB, IL-6/JAK/STAT3, p53, apoptosis (red box), PI3K/AKT/mTOR, inflammatory response (red box). Module 2 (M2, orange bars): associated pathways including KRAS, complement, cholesterol homeostasis, reactive oxygen species, and coagulation.

Conversely, several proteins with key homeostatic and pro-angiogenic roles were significantly depleted in CSE-derived EVs. Most notably, Milk Fat Globule EGF Factor 8 (MFG-E8/Lactadherin), a glycoprotein critically involved in efferocytosis, angiogenesis, and vascular homeostasis, was markedly downregulated. Similarly, Ephrin-B2 (EFNB2), a regulator of vascular development, and VPS4B, an ESCRT pathway component central to EV biogenesis itself, were significantly reduced. Losses in Kinesin Family Member 2B (KIF2B), Lipocalin-1 (LCN1), and protein phosphatase PPP1CA further reflected disruption of intracellular transport, lipid metabolism, and signaling painting a picture of trophoblasts under duress, shedding EVs that carry signatures of injury while losing protective and regenerative capacity. Taken together, these changes suggest that oxidative stress fundamentally reshapes the trophoblast EV proteome enriching for damage-associated cargo while stripping away protective and pro-angiogenic factors.

Notably, PLAP-expressing CTCs exhibited molecular alterations comparable to those previously reported in placental trophoblast cells, indicating that the fetal membrane compartment itself manifests PE-like injury signatures. This finding suggests that fetal membrane derived trophoblasts and their EVs may contribute directly to PE pathophysiology and may represent an additional target for therapeutic intervention.

IPA functional analysis predicted increased apoptosis based on the network of CSE-altered proteins (Figure 3B). The apoptosis network incorporated multiple convergent pro-death signals, consistent with CSE-induced trophoblast injury. Simultaneously, IPA predicted reduced angiogenesis following CSE exposure (Figure 3C). The anti-angiogenic network included downregulation of pro-vascular proteins and loss of growth factor signaling, indicating that CSE-damaged CTC-EVs carry cargo that suppresses vascular function at the feto-maternal interface. Consistent with oxidative injury, CSE exposure strongly activated apoptotic programs, reflected by increased abundance of CASP3, BAX, and multiple p53-regulated stress-response proteins. Proteomic and network-level analyses of CTC-derived EVs revealed that CSE exposure activated TNF-driven apoptotic signaling and p53-dependent cellular stress pathways (Supplementary Figure1 A, B) aligning with established mechanisms of PE-related trophoblast injury. In parallel, CSE markedly suppressed angiogenic and ECM-remodeling proteins, including VEGFA, MMP14, and COL4A1 indicating loss of endothelial support and diminished tissue remodeling capacity. This molecular pattern closely parallels the angiogenic insufficiency and impaired vascular remodeling characteristic of PE-affected placental and fetal membrane tissues.

To move beyond individual protein changes and understand the broader biological programs at play, we performed pathway enrichment analysis on differentially expressed proteins between CSE-treated and control CTC-derived sEVs. Hierarchical clustering revealed two major coordinated expression modules (Figure 3D).

The first module (M1), representing pathways activated under CSE exposure, was dominated by inflammatory and cell death signaling. TNFα signaling via NF-κB and IL-6/JAK/STAT3 signaling featured prominently, pointing to a distinct pro-inflammatory program embedded within the EV cargo. Alongside these, activation of the p53 pathway and apoptotic signaling indicated that CSE-challenged trophoblasts were not merely inflamed but actively primed for cell death, a finding consistent with the tissue injury phenotype observed in preeclampsia. Interferon-γ response, PI3K/AKT/mTOR signaling, and hypoxia response pathways were also enriched, reflecting the convergence of immune activation, survival signaling, and oxygen-sensing mechanisms that typify the stressed placental microenvironment. Notably, enrichment of allograft rejection and TGF-β signaling pathways hinted at disrupted immune tolerance at the feto-maternal interface, a hallmark of preeclampsia pathogenesis.

The second module (M2) captured a set of contextual metabolic and homeostatic pathways that, while not classically inflammatory, are increasingly recognized as contributors to placental dysfunction. KRAS signaling and interferon-α response suggested altered proliferative and innate immune signaling, while enrichment of cholesterol homeostasis, fatty acid metabolism, and heme metabolism pointed to broader metabolic rewiring under oxidative stress. Particularly striking was the enrichment of the reactive oxygen species pathway and complement cascade both of which directly reinforce the oxidative and immunological damage initiated by CSE. Coagulation pathway enrichment in M2 aligned with the striking upregulation of Factor X (F10) observed at the individual protein level, reinforcing a coherent pro-thrombotic EV signature.

Taken together, these two modules paint a picture of trophoblast derived EVs that carry not just isolated stress markers but an integrated, multi-layered program of inflammation, apoptosis, metabolic disruption, and coagulation effectively serving as molecular messengers of fetal membrane distress and oxidative stress (Table 3).

**Table 3:**
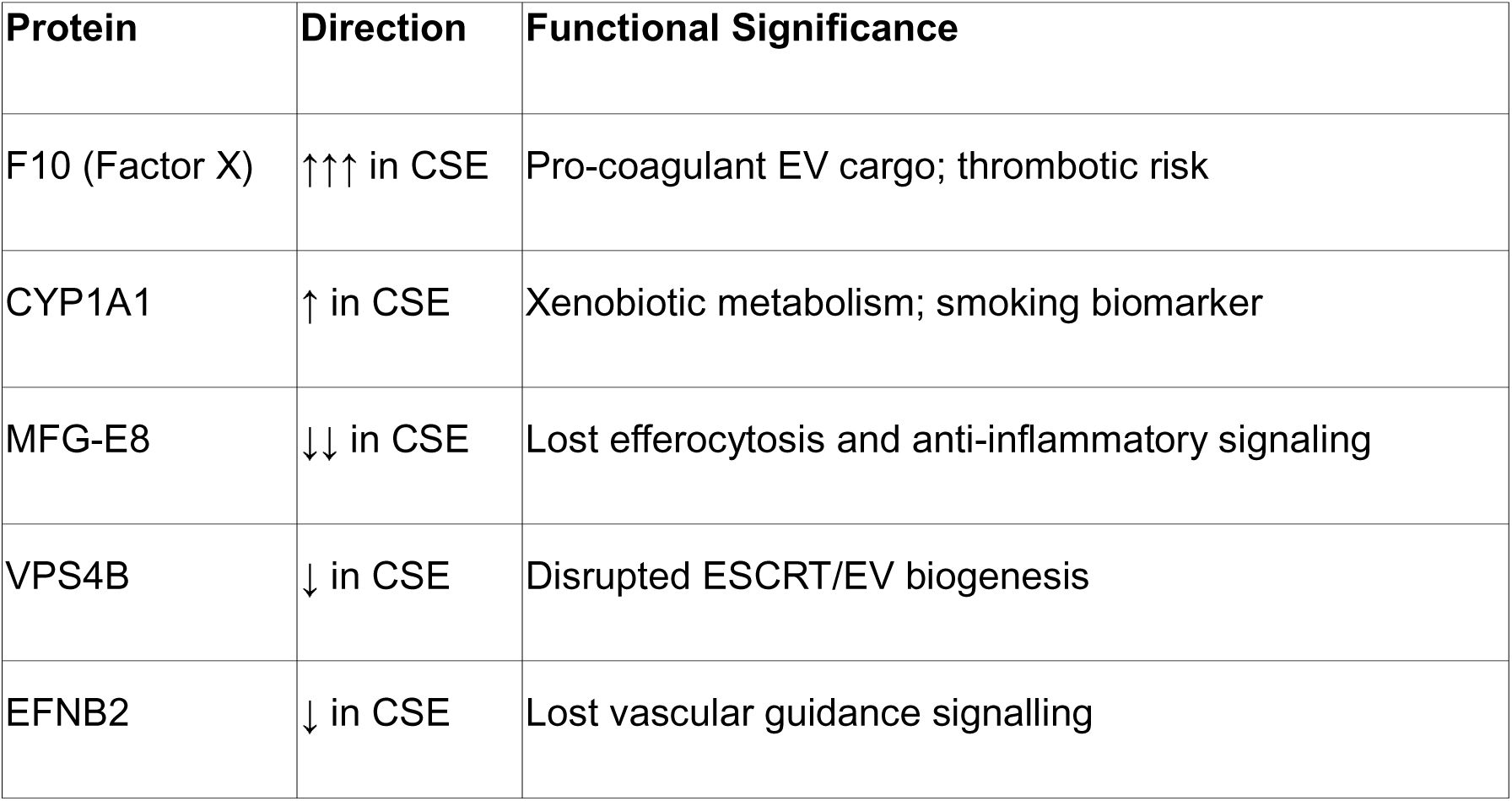
CSE-Induced damage signature proteins.

### 3.4 Prophylactic Aspirin co-treatment reduces apoptosis in a dose-dependent manner and restores angiogenesis exclusively at low dose

#### Low-dose co-treatment (Asp4-CSE vs. CSE)

As angiogenesis impairment is a major pathology associated with PE, we first examined how prophylactic aspirin influences angiogenic signaling within CTCs, using EV cargo as a proxy for underlying cellular processes.

Under prophylactic conditions, CTC-derived EVs exhibited moderate recovery of angiogenic and structural remodeling pathways. Volcano plot analysis revealed distinct proteomic reprogramming (Figure 4A), with upregulation of SLC4A7, LGR4 (Wnt signalling receptor; endothelial proliferation), GPRC5C, IL17RA (immune modulation), THY1/CD90 (angiogenesis), and RHBIDF2. IPA predicted reduced apoptosis (Figure 4C), with a network involving B2M, YBX1, CAPNS1, RPS27A, DDX21, RAD23B, FUBP1, and MCAM. Critically, IPA simultaneously predicted increased angiogenesis (Figure 4D), with a network incorporating ANXA3, CAPN1, CAV1 (caveolin-1; VEGF signalling scaffold), COLTA1, SERPINC1, SARS1, RHOA, MMP14 (MT1-MMP; matrix remodeling), MYOF (myoferlin; VEGFR2 signalling), and NF1.

**Figure 4.**
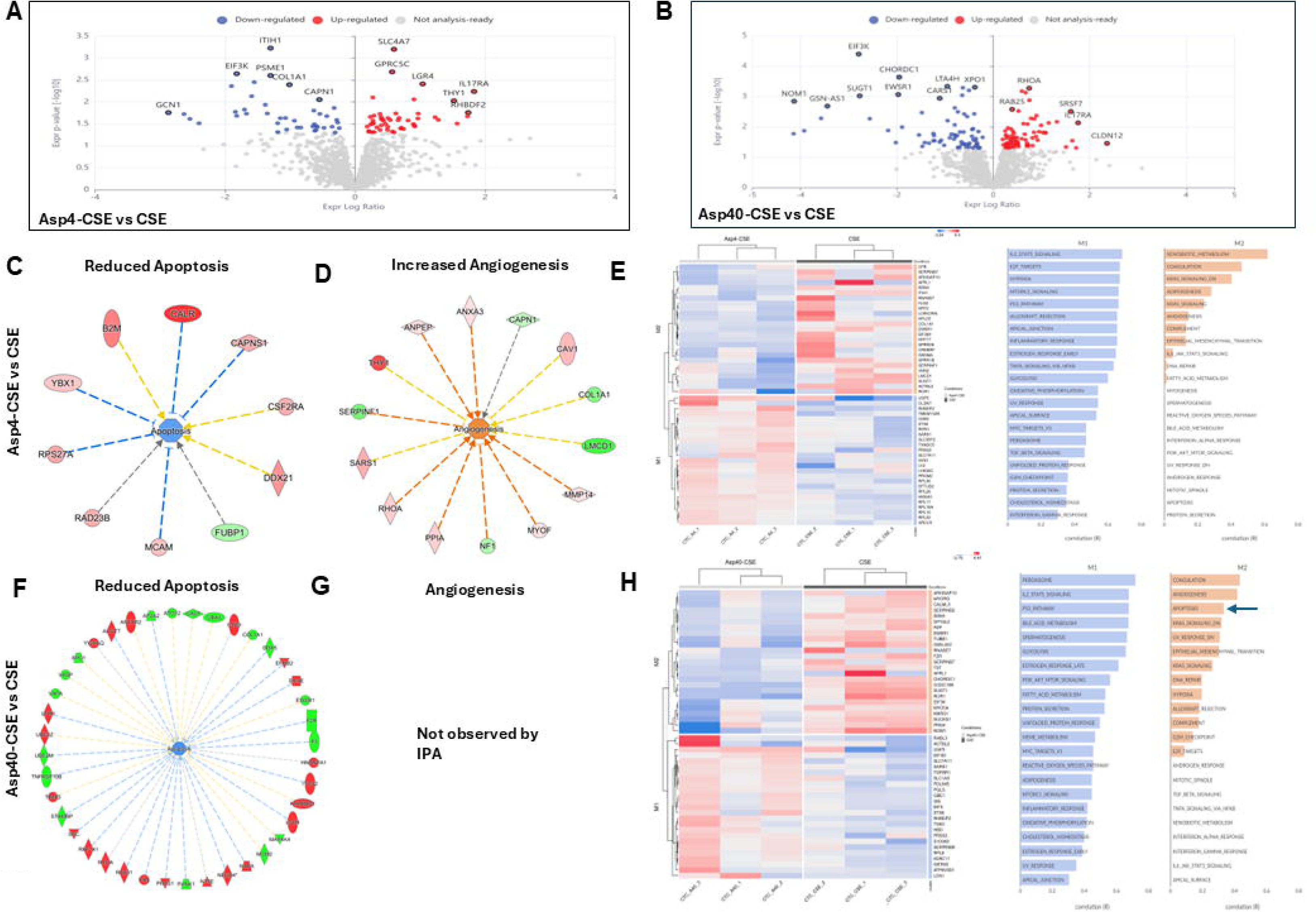
Prophylactic aspirin co-treatment reprograms CSE-induced trophoblast EV protein expression in a dose-dependent manner. (A) Volcano plot: Asp4-CSE vs. CSE. Significantly upregulated proteins include SLC4A7, LGR4, GPRC5C, IL17RA, THY1, and RHBIDF2. (B) Volcano plot: Asp40-CSE vs. CSE. Fewer significant changes, including CHORDC1, XPNPEB2, LTA4H, RHOA, and CLDN1. (C) IPA network predicting reduced apoptosis in Asp4-CSE vs. CSE, involving B2M, YBX1, CAPNS1, DDX21, RAD23B, FUBP1, and MCAM. (D) IPA network predicting increased angiogenesis in Asp4-CSE vs. CSE, involving ANXA3, CAPN1, CAV1, SERPINC1, RHOA, MMP14, MYOF, and NF1. (E) Hierarchical clustering heatmap and pathway enrichment for Asp4-CSE vs. CSE. Blue arrow indicates angiogenesis enrichment. (F) IPA network predicting reduced apoptosis in Asp40-CSE vs. CSE. (G) Angiogenesis prediction for Asp40-CSE vs. CSE: “Not observed by IPA.” (H) Heatmap and pathway enrichment for Asp40-CSE vs. CSE, with Angiogenesis and apoptosis enrichment highlighted (red boxes). Each volcano plot point represents a protein plotted by log_₂_ expression ratio (x-axis) and −log_₁₀_ adjusted *p*-value (y-axis).

Asp4-CSE-Asp4; (prophylaxis-exposure-treatment) enhanced expression of VEGFA (log_₂_ FC +0.7), COL4A1 (log_₂_ FC +0.8, p = 0.004), and MMP14 (p = 0.047), suggesting partial restoration of endothelial signaling and extracellular matrix (ECM) organization, indicative of the membrane remodeling process after oxidative stress-induced damage (Fig. 5A, Fig 5 C,D,E). Functional enrichment analysis confirmed the activation of VEGF, integrin, and cholesterol metabolism pathways after prophylactic Asp treatment.

**Figure 5:**
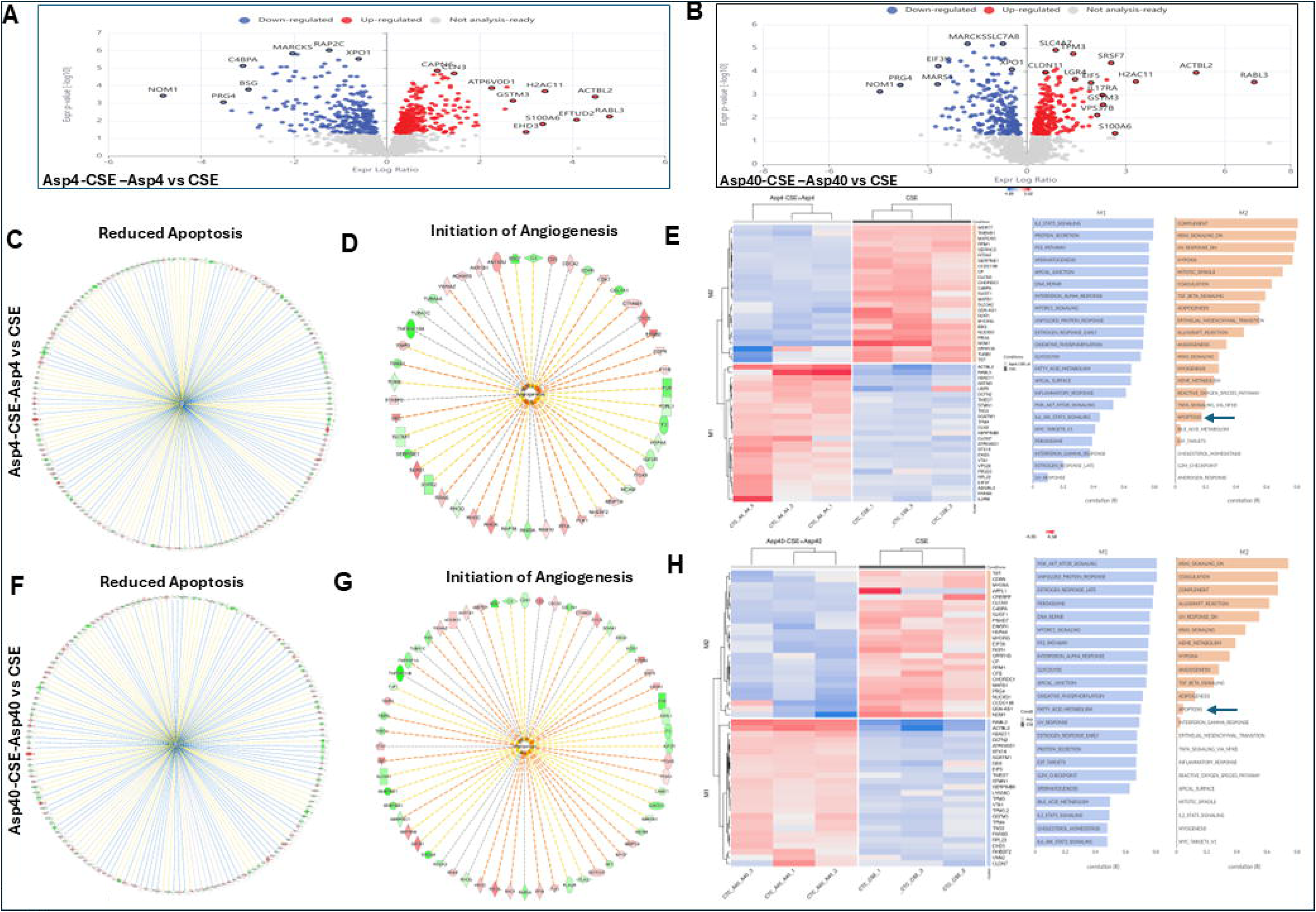
L**o**w**-dose prophylactic aspirin initiates to restore angiogenic signaling suppressed by oxidative injury**. (A–B) Volcano plots depict differential gene expression profiles under prophylactic aspirin treatment paradigms in CTCs exposed to CSE. (C–E) Asp4-CSE-Asp4 vs CSE: Low-dose aspirin pretreatment reverses CSE-induced suppression, up-regulating SLC4A7, LGR4, and RHOJ, and activating pathways associated with endothelial migration and extracellular matrix remodeling, consistent with enhanced angiogenesis.(F–H) Asp40-CSE-Asp40 vs CSE: High-dose aspirin exerts minimal or neutral effects on angiogenic pathways, with only modest transcriptional changes and absence of significant enrichment in angiogenesis-related gene sets. Volcano plots show differential expression (log_₂_ fold-change vs –log_₁₀_ p-value); heatmaps and pathway enrichment plots illustrate relative gene expression and pathway correlation across treatment conditions.

#### High-Dose Co-Treatment (Asp40-CSE vs. CSE)

High-dose co-treatment produced fewer significant changes (Figure 4B), with modest alterations in CHORDC1, XPNPEB2, LTA4H, SRSF4, RHOA, and CLDN1. IPA predicted reduced apoptosis (Figure 4F) but critically, angiogenesis was not predicted by IPA noted as “Not observed by IPA” (Figure 4G). Pathway enrichment showed a distinct profile with coagulation-related pathway enrichment (Figure 4H).

Low-dose prophylactic aspirin achieves dual rescue (anti-apoptotic + pro-angiogenic), while high-dose prophylaxis prevents apoptosis but fails to restore angiogenic signaling. This dose-dependent dissociation is a principal finding of the study (Summarized in Table 4).

**Table 4:**
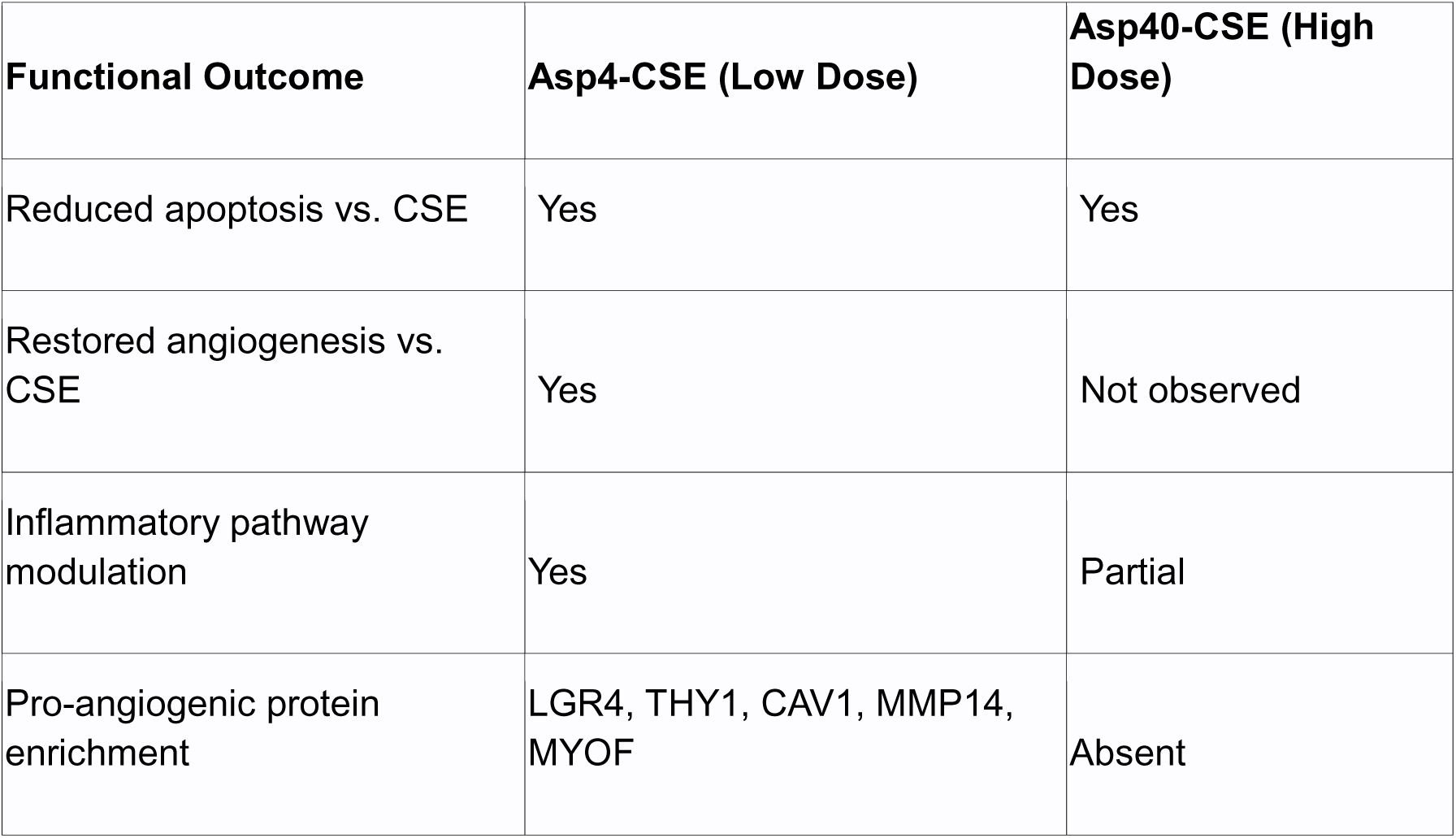
Functional outcome of prophylactic low or high dose of Aspirin.

### 3.5 Continuous low-dose prophylactic aspirin initiate angiogenic signaling suppressed by oxidative injury

To evaluate whether extending aspirin exposure (before AND after CSE) provides additional benefit, we analysed the combined treatment groups Asp4-CSE-Asp4 and Asp40-CSE-Asp40 compared with CSE alone (Figure 5).

#### Low-dose continuous (Asp4-CSE-Asp4 vs. CSE)

Extended low-dose aspirin exposure produced reprogramming (Figure 5A), with upregulation of MARCKS (membrane dynamics), RAP2C, CAPNS3, ATPV0D1 (vacuolar ATPase; EV biogenesis), H2AC11, GSTM3 (glutathione S-transferase; antioxidant defense), S100A6, EFTUD2, and RABL3 (vesicular trafficking). IPA predicted both reduced apoptosis (Figure 5C) and initiation of angiogenesis (Figure 5D). Pathway enrichment confirmed restoration of angiogenesis-related signaling (Figure 5E, blue arrows).

#### High-dose continuous (Asp40-CSE-Asp40 vs. CSE)

Unlike high-dose co-treatment alone, the combined high-dose condition did show initiation of angiogenesis (Figure 5G), alongside reduced apoptosis (Figure 5E, F). Pathway enrichment showed TGF-β and angiogenesis-related pathway restoration (Figure 5G, H).

This demonstrates that extending aspirin exposure (before and after CSE) partially compensates for the dose-dependent limitation, with the most restoration in the low-dose combined group (summarized in Table 5).

**Table 5:** Functional outcome of continuous prophylactic low or high dose of Aspirin.

| **Functional Outcome** | **Asp4-CSE-Asp4 (Low Continuous)** | **Asp40-CSE-Asp40 (High Continuous)** |
| --- | --- | --- |
| Reduced apoptosis vs. CSE | Yes | Yes |
| Initiation of angiogenesis | Yes (Strong) | Yes (partial) |
| Antioxidant pathway enrichment | GSTM3 upregulated | Less prominent |
| EV biogenesis markers | ATPV0D1 upregulated | Less clear |
| Vesicular trafficking | RABL3, RAP2C | MARCKS, MARCKSL1 |

### 3.6 Prophylactic aspirin suppress apoptosis and inflammation in CTC-EVs

To determine whether prophylactic aspirin modulates CSE-induced inflammation, IPA network analysis was performed specifically for inflammatory response pathways (Figure 6).

**Figure 6.**
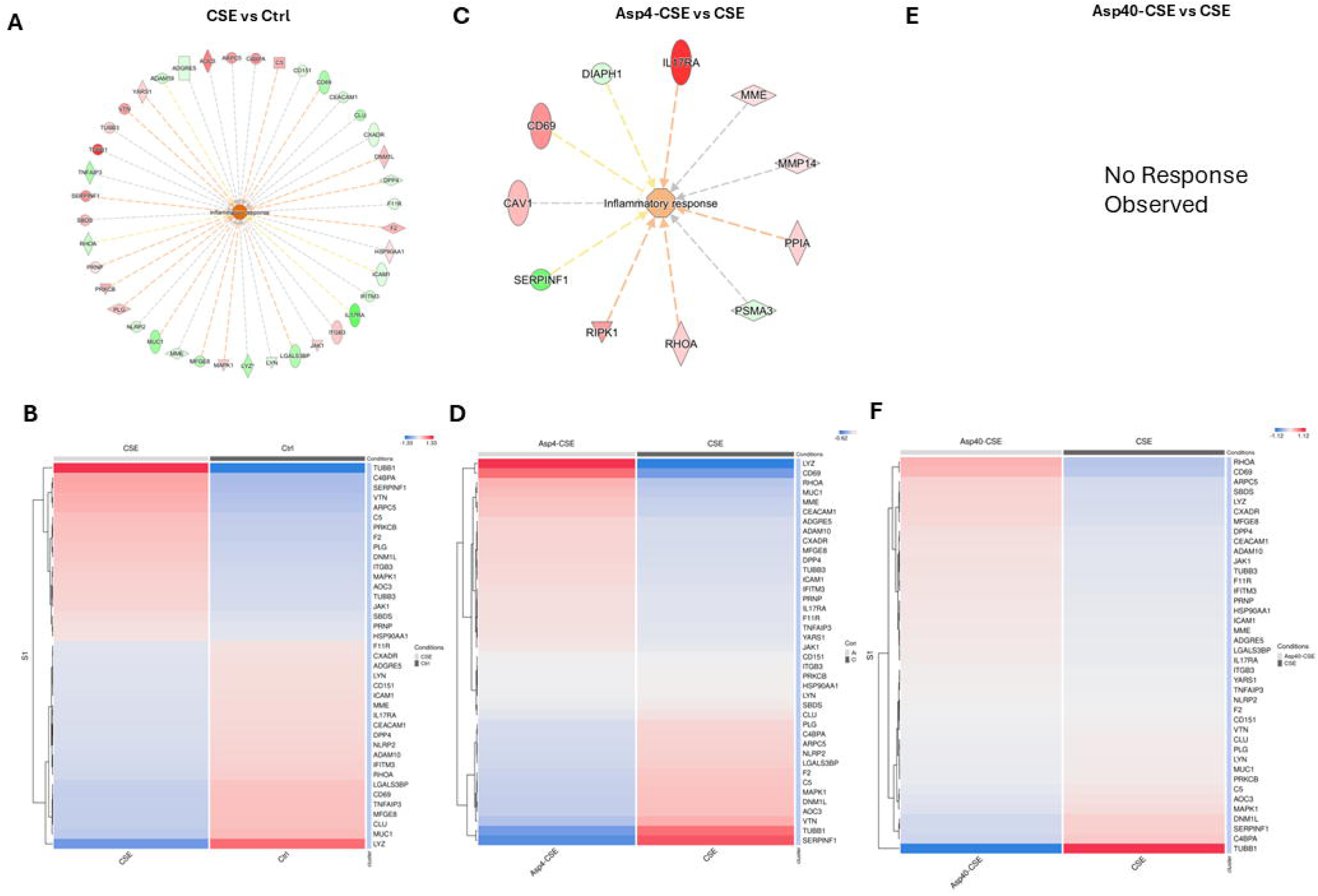
**Timing and dose constrain aspirin-mediated modulation of inflammatory transcriptional networks**. (A) IPA inflammatory response network: CSE vs. Control. CSE activates a broad inflammatory network involving C5, CXADR, TNF, JAK1, ITGB3, and multiple adhesion molecules. (B) Hierarchical heatmap of inflammation-associated EV proteins, CSE vs. Control. (C) IPA inflammatory response network: Asp4-CSE vs. CSE. Low-dose prophylactic aspirin produces a targeted network involving IL17RA, CD69, DIAPH1, CAV1, MMP14, SERPINF1, RIPK1, RHOA, and PPIA. (D) Heatmap: Asp4-CSE vs. CSE, showing normalization of inflammatory gene expression. (E) IPA inflammatory network: Asp40-CSE vs. CSE: “No Response Observed.” (F) Heatmap: Asp40-CSE vs. CSE, showing minimal transcriptional reprogramming. Inflammatory modulation is exclusive to low-dose prophylactic aspirin and is absent at high dose.

#### CSE vs. Control

(Figure 6A–B). CSE activated a broad inflammatory signaling network involving C5, CKADR, FKBP1A, C10TNF3, TRIM25, TNF, THBS4, THBS1, PLAU, LGALS3BP, JAK1, ITGB3, IGKC, IGHG1, HSPG2, and GCLM, with coordinate upregulation of adhesion molecules (ICAM1, CEACAM1, CD151), proteases (ADAM10, DPP4, MME), signaling kinases (LYN, MAPK1, JAK1), and inflammatory mediators. Hierarchical clustering confirmed clear separation of CSE and control samples across inflammation-associated genes (Figure 6B), with proteins including TUBB1, C4BPA, SERPINF1, VTN, ARPC5, and MFG-E8 among the most differentially expressed.

#### Asp4-CSE vs. CSE (Figure 6C–D)

Low-dose prophylactic aspirin produced a targeted inflammatory modulation network centered on the inflammatory response node, involving IL17RA (upregulated), DIAPH1, CD69 (upregulated), MME, CAV1, MMP14, SERPINF1 (downregulated), RIPK1, RHOA, PPIA, and PSMA3. The heatmap (Figure 6D) showed a clear shift toward normalization of CSE-induced inflammatory gene expression, with notable restoration of MFG-E8 expression and modulation of CEACAM1, ADAM10, and ICAM1.

#### Asp40-CSE vs. CSE (Figure 6E–F)

In striking contrast, high-dose prophylactic aspirin produced “No Response Observed” for inflammatory networks. IPA did not detect coordinated inflammatory response modulation. The heatmap (Figure 6F) showed minimal transcriptional reprogramming compared with the low-dose condition.

This finding demonstrates that inflammatory modulation by prophylactic aspirin is exclusive to low dose and is absent at high dose, paralleling the angiogenesis dose-dependency.

### 3.7 Continuous Aspirin followed by CSE induced oxidative stress attenuates inflammation and restores metabolic homeostasis

To assess whether extending aspirin after CSE exposure enhances inflammatory resolution, combined treatment groups were analyzed for inflammatory networks (Figure 7).

**Figure 7.**
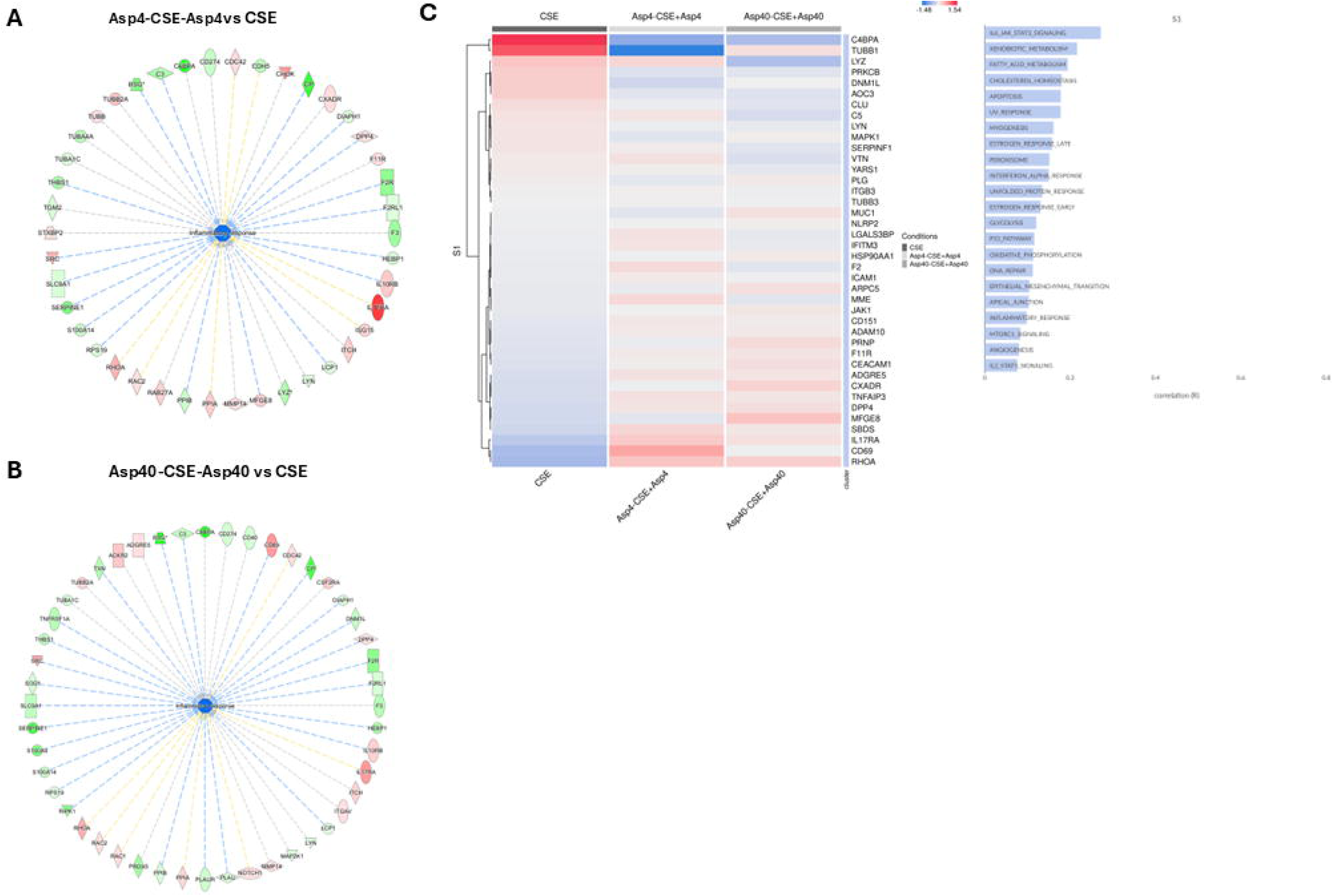
Aspirin continuation following oxidative stress selectively attenuates inflammatory signaling and restores metabolic homeostasis. (A) IPA inflammatory response network: Asp4-CSE-Asp4 vs. CSE. Coordinated downregulation (green nodes) of inflammatory mediators with mixed activation/inhibition relationships. (B) IPA inflammatory network: Asp40-CSE-Asp40 vs. CSE. Broader but less directional transcriptional response. (C) Hierarchical heatmap of inflammation-associated proteins across CSE, Asp4-CSE-Asp4, and Asp40-CSE-Asp40 conditions. Asp4-CSE-Asp4 shows the most consistent normalization. Key proteins: C4BPA, TUBB1, LYZ, PRKCB, CLU, MAPK1, SERPINF1, VTN, MFG-E8, ICAM1, CD151, CD69, RHOA. S1 module pathway correlations showing aspirin continuation influences IL-6/JAK-STAT signaling, xenobiotic metabolism, cholesterol homeostasis, oxidative phosphorylation, DNA repair, apical junction, inflammatory response, angiogenesis, and mTORC1 signaling. Node colour: red = upregulated; green = downregulated. Edge colour: orange = predicted activation; blue = predicted inhibition.

Asp4-CSE-Asp4 vs. CSE (Figure 7A). IPA revealed a large, coordinated network showing predominant downregulation (green nodes) of inflammatory mediators, with both activation (orange edges) and inhibition (blue edges) relationships, indicating active remodeling of the inflammatory landscape. The network included CXADR, DPP4, F2K, F11R, CD274, CXCL2, ADAM10, and numerous downstream targets.

Asp40-CSE-Asp40 vs. CSE (Figure 7B). The high-dose combined condition also showed inflammatory network modulation, though with a broader but less directional pattern featuring both up-and downregulated nodes.

Heatmap comparison (Figure 7C). Hierarchical clustering of inflammation-associated proteins across CSE, Asp4-CSE-Asp4, and Asp40-CSE-Asp40 conditions demonstrated that Asp4-CSE-Asp4 showed the most consistent normalization toward control-like expression, while Asp40-CSE-Asp40 showed partial modulation. Key normalized proteins included C4BPA, TUBB1, LYZ, PRKCB, DMKL1, AOC3, CLU, LYN, C5, MAPK1, SERPINF1, VTN, and MFG-E8.

Pathway correlations (Figure 7D). The S1 module pathway analysis revealed that continuous aspirin treatment influenced IL-6/JAK-STAT signalling, xenobiotic metabolism, fatty acid metabolism, cholesterol homeostasis, oxidative phosphorylation, DNA repair, apical junction integrity, inflammatory response, angiogenesis, and mTORC1 signaling.

These data indicate that continued low-dose aspirin following oxidative insult preferentially suppresses inflammatory signaling while simultaneously promoting metabolic and epithelial homeostasis

### 3.8 EVs encode partial attenuation of apoptotic and necrotic signaling following post-injury therapeutic aspirin exposure

**CSE Damage in the Therapeutic Arm (CSE vs. Control; Figure 8A-D):**In the therapeutic experimental arm, CSE exposure (designated CSE 4, reflecting the therapeutic protocol timing) induced significant proteomic changes, with upregulation of SLCAT11, OGLN, and stress-responsive proteins (partial stress adaptation), and downregulation of PLSCR1, GUL8, LGALS3BP, TSPAN6, and KRT66. IPA predicted increased apoptosis (Figure 8B), increased necrosis (Figure 8C), and activated protein regulatory networks (Figure 8D same as supplementary figure 3A). This indicates an EV-encoded shift away from pro-apoptotic network states rather than complete suppression of death signaling (F

**Figure 8.**
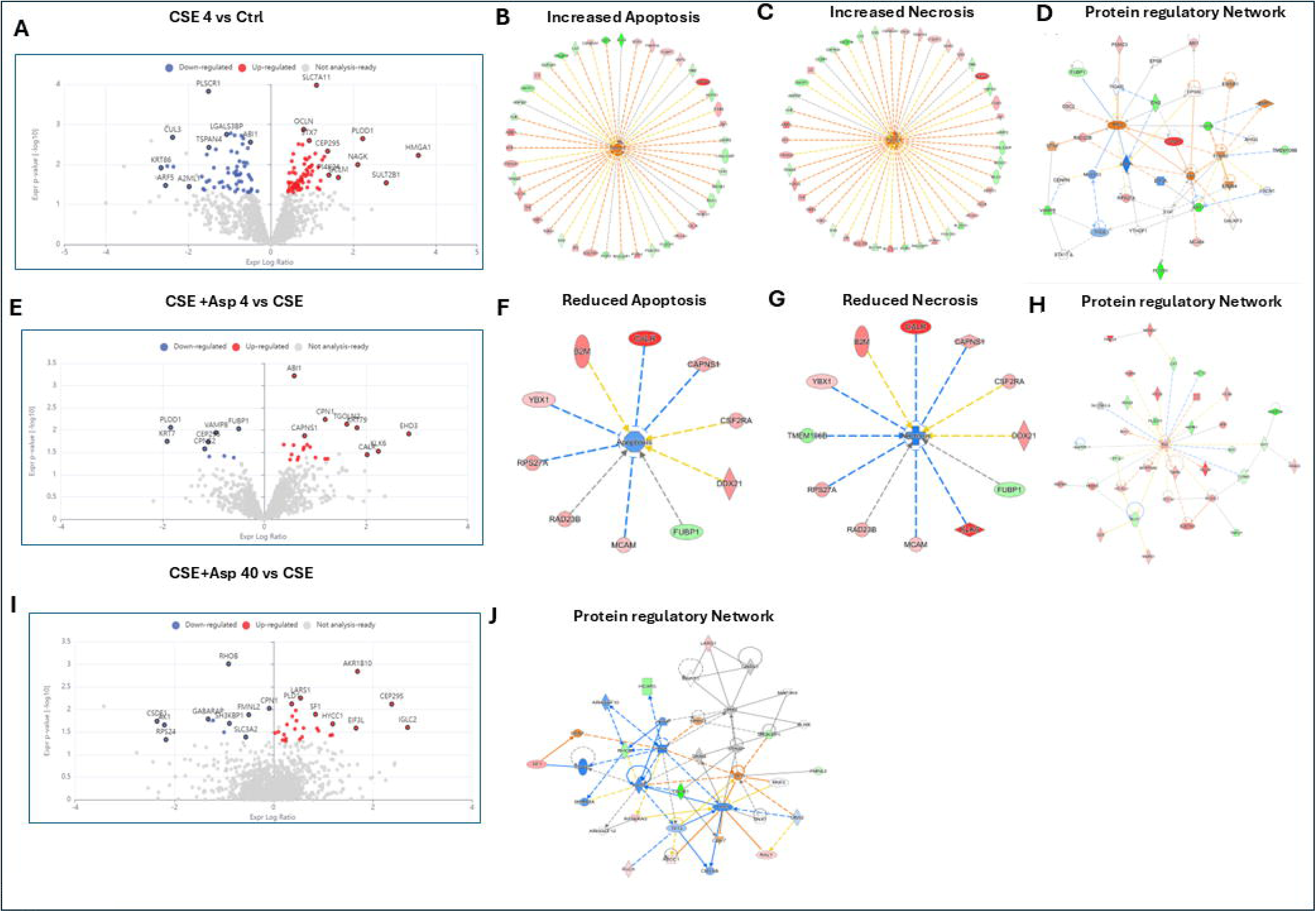
Therapeutic aspirin attenuates oxidative stress–induced apoptotic and necrotic signalling in CTC-EVs. (A) Volcano plot: CSE vs. Control (therapeutic arm). Upregulated: SLC7A11, OGLN, CEP205, NAGK, HMAGA1, SULTZ81. Downregulated: PLSCR1, GUL8, LGALS3BP, TSPAN6, KRT66. (B–C) IPA networks predicting increased apoptosis (B) and increased necrosis (C) in CSE vs. Control. (D) Protein regulatory network: CSE vs. Control. (E) Volcano plot: CSE+Asp4 vs. CSE. Fewer significant changes; notable: AB1, PLOD1, CAPNS1, EHQ2. (F–G) IPA networks predicting reduced apoptosis (F) and reduced necrosis (G) in CSE+Asp4 vs. CSE. Networks involve CAPNS1, YBX1, CSF2RA, DDX21, RPS27A/K, RAD23B, FUBP1, MCAM. (H) Protein regulatory network: CSE+Asp4 vs. CSE. (I) Volcano plot: CSE+Asp40 vs. CSE. Broader responses include RHO5, AKR1B10, CEP295, SF1, IGLC2. (J) Protein regulatory network: CSE+Asp40 vs. CSE, showing extensive remodeling.

#### Low-Dose Therapeutic Aspirin (CSE+Asp4 vs. CSE; Figure 8E–H)

Therapeutic low-dose aspirin produced fewer differentially expressed proteins than prophylactic conditions (Figure 8E), with modest changes in AB1, PLOD1, VAVHE8, CAPNS1, IGOLM75, EHQ2, and CALA55. Despite limited individual protein changes, IPA predicted reduced apoptosis (Figure 8F), with a network involving GAPR, CAPNS1, YBX1, CSF2RA, DDX21, RPS27K, RAD23B, FUBP1, and MCAM, and reduced necrosis (Figure 8G), with a parallel network including B2K, CAPNS1, YBX1, TMEM198B, CSF2RA, DDX21, RPS27A, RAD23B,

FUBP1, and MCAM. EVs generated under low-dose therapeutic aspirin exposure also displayed reduced enrichment of inflammatory and necrosis-associated pathways (Fig. 8E–H, Supplementary Figure 3A).

#### High-Dose Therapeutic Aspirin (CSE+Asp40 vs. CSE; Figure 8I–J)

High-dose therapeutic aspirin (Figure 8I) produced a broader transcriptional response than low-dose, with upregulation of RHO5, AKR1B10, CEP295, SF1, IGLC2, and others. The protein regulatory network (Figure 8J) demonstrated extensive remodeling with both activation and inhibition interactions. In contrast to low dose, EV cargo from high-dose aspirin conditions showed decreased representation of survival-associated proteins, including MCL1-and AKT1-linked networks, suggesting that excessive pharmacologic inhibition may constrain EV-encoded adaptive recovery programs (Fig. 8I–J). Network-level analyses of EV proteomic states demonstrated (Supplementary Fig 1A, B), that low-dose therapeutic aspirin most effectively limited EV-encoded apoptotic signaling while preserving reparative programs (Supplementary Fig 3A-B), whereas high-dose exposure (Supplementary Fig 3C-D) broadly suppressed both damage and repair-associated EV networks, consistent with a dormant or constrained adaptive state. Both therapeutic doses attenuate CSE-induced apoptosis and necrosis, but through networks of modest individual protein changes, suggesting that therapeutic aspirin achieves partial rescue through subtle, distributed effects rather than the reprogramming seen with prophylactic treatment.

### 3.9 Therapeutic Aspirin fails to reverse CSE-induced inflammatory programming and timing-dependent prophylactic effects reveal limited capacity of aspirin to reprogram CTC inflammation

Exposure to CSE elicited a strong inflammatory response in CTCs, marked by pronounced increases in IL-6 and IL-8 secretion, minimal alteration in TNF, and a modest decline in IL-10 supported by IPA network analysis (Fig 9A-D). In the therapeutic treatment paradigm, aspirin did not mitigate this CSE-driven activation. Cytokine levels in the CSE+Asp4 and CSE+Asp40 groups remained similar to, or in some cases exceeded, those observed with CSE alone. No Networks were produced by IPA in case of treatment (Fig 9 B, C) TNF production was largely unaffected. Notably, IL-10 concentrations were further reduced by aspirin, leading to elevated IL-6/IL-10 and TNF/IL-10 ratios. Consistent with these patterns, the composite z-scored Inflammation Index (IL-6 + IL-8 + TNF − IL-10) reached its highest value in the CSE+Asp40 condition, indicating that therapeutic aspirin failed to resolve and may even intensify the inflammatory imbalance in CTCs shown in figure 9A. In the prophylactic paradigm, similar resistance to aspirin was observed. Co-treatment with Asp4 or Asp40 failed to prevent CSE-induced IL-6 and IL-8 up-regulation, and these groups displayed higher IL-6/IL-10 ratios than CSE alone. Only the continuous exposure schedule (Asp→CSE→Asp) partially restored IL-10 and produced a modest reduction in the Inflammation Index shown in Figure 9D, E, suggesting that transient prophylactic dosing is insufficient to reprogram CTC inflammatory output.

**Figure 9.**
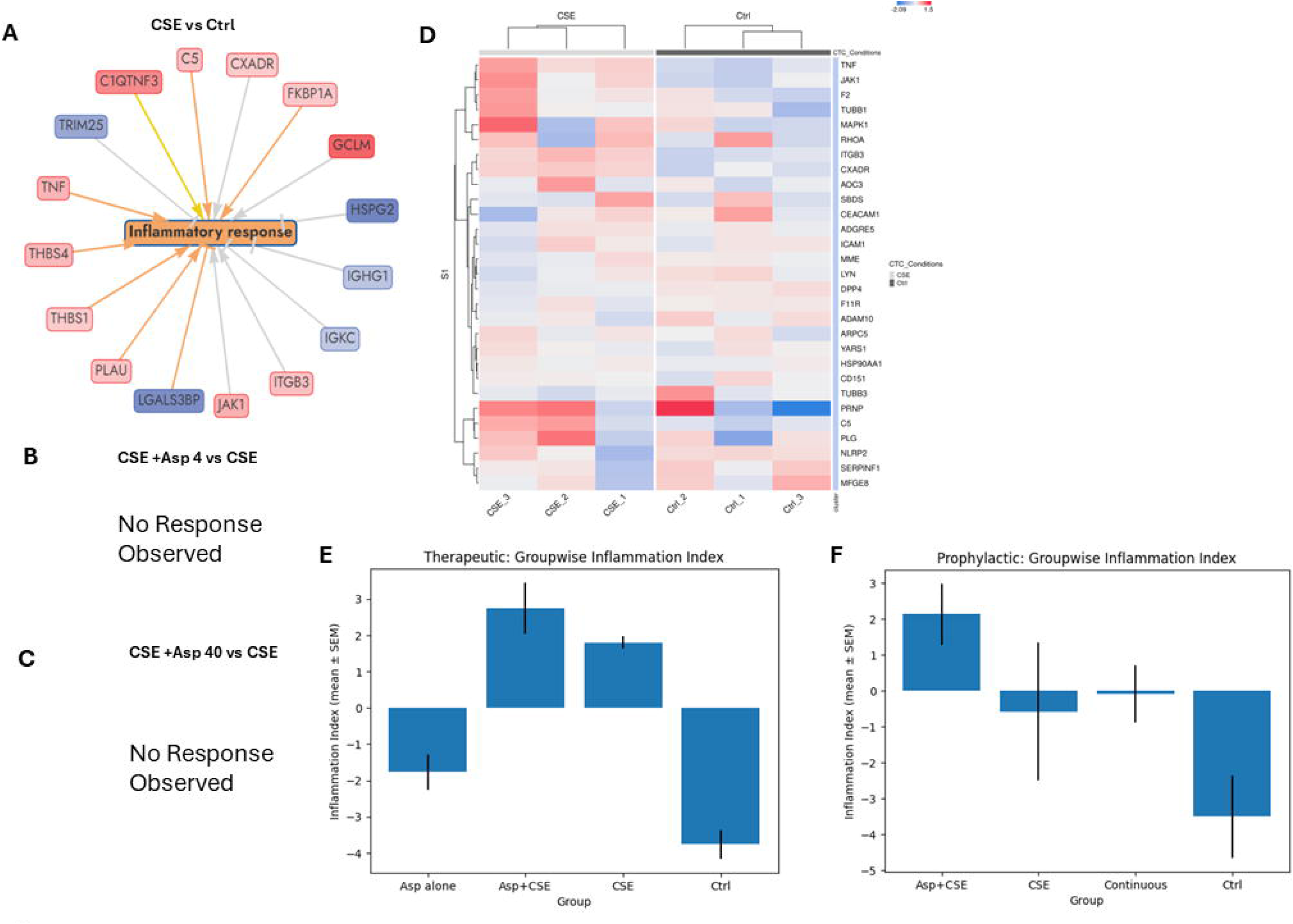
CSE induces a strong inflammatory transcriptional program that is not reversed by therapeutic aspirin and composite inflammation index reveals that only continuous aspirin normalizes inflammatory signaling. (A) IPA inflammatory response network: CSE vs. Control. CSE induces coordinated upregulation of inflammatory mediators including C5, CKADR, FKBP1A, GCLM, HSPG2, IGHG1, TRIM25, TNF, THBS4/THBS1, PLAU, JAK1, ITGB3. (B) CSE+Asp4 vs. CSE: “No Response Observed” therapeutic low-dose aspirin does not produce a detectable inflammatory network response. (C) CSE+Asp40 vs. CSE: “No Response Observed.” (D) Hierarchical heatmap of inflammation-associated proteins, CSE vs. Control. CSE samples cluster distinctly from controls, with consistent induction of TNF, JAK1, F2, TUBB1, MAPK1, RHOA, ITGB3, CXADR, and suppression of PRNP, TUBS3, NLRP2, SERPINF1, MFG-E8. Together, these data demonstrate that therapeutic aspirin is insufficient to reverse CSE-established inflammatory transcriptional states at either dose, highlighting the critical importance of aspirin timing in immunomodulation at the feto-maternal interface. (E) Therapeutic arm: Groupwise Inflammation Index calculated as z(IL-6) + z(IL-8) + z(TNF) − z(IL-10). Aspirin alone shows low inflammation comparable to control (Ctrl). CSE elevates the index. Therapeutic aspirin following CSE (Asp+CSE) does not reduce the index and trends higher than CSE alone, indicating failure to resolve and potential exacerbation of inflammatory signaling. (F) Prophylactic arm: Asp+CSE (co-treatment) shows an elevated Inflammation Index higher than CSE alone. However, the Continuous schedule (Asp-CSE-Asp; red box) preserves the Inflammation Index near baseline, with values comparable to control. CSE alone shows a modest positive index. Data represent mean ± SEM of n = 3 biological replicates. These data demonstrate that only continuous aspirin exposure (before and after CSE) achieves normalization of the inflammatory milieu; neither single-phase prophylactic nor therapeutic aspirin is sufficient.

### 3.10 Comparative analysis across all treatment paradigms

Integrating findings from all prophylactic and therapeutic treatment, a hierarchy of angiogenic rescue emerges summarized in Table 6. The critical finding is that angiogenic rescue is preferentially achieved by low-dose aspirin, with the most distinct restoration occurring in the Asp4-CSE co-treatment and Asp4-CSE-Asp4 combined conditions. High-dose aspirin co-treatment (Asp40-CSE) fails to rescue angiogenesis entirely, while extended high-dose exposure (Asp40-CSE-Asp40) partially restores it. The pro-angiogenic program restored by low-dose aspirin involves a coordinated set of proteins with established roles in vascular development: the key protein drivers are listed in table 7

**Table 6:** Comparative analysis of prophylactic and therapeutic treatment.

| **Treatment Paradigm** | **Apoptosis Rescue** | **Angiogenesis Rescue** | **Overall Assessment** |
| --- | --- | --- | --- |
| CSE alone | Increased apoptosis | Reduced angiogenesis | Damage baseline |
| Asp4-CSE (Prophylactic low) | Reduced | Increased | Best co-treatment |
| Asp40-CSE (Prophylactic high) | Reduced | Not observed | Partial protection |
| Asp4-CSE-Asp4 (Continuous low) | Reduced | Initiated | Best overall |
| Asp40-CSE-Asp40 (Continuous high) | Reduced | Initiated (partial) | Good protection |

**Table 7:** Proangiogenic protein regulators observed in this study.

| Protein | Function | Condition Observed |
| --- | --- | --- |
| LGR4 | Wnt signalling receptor; promotes endothelial proliferation | Asp4-CSE ↑ |
| THY1 (CD90) | Cell surface glycoprotein; angiogenesis and adhesion | Asp4-CSE ↑ |
| CAV1 (Caveolin-1) | Membrane scaffolding; VEGF signalling | Asp4-CSE ↑ |
| MMP14 (MT1-MMP) | Extracellular matrix remodelling; invasion | Asp4-CSE ↑ |
| MYOF (Myoferlin) | Membrane repair; VEGFR2 signalling; angiogenesis | Asp4-CSE ↑ |
| RHOA | Rho GTPase; endothelial migration | Asp4-CSE ↑, Asp40-CSE ↑ |
| SLC4A7 | Bicarbonate transporter; pH regulation in angiogenesis | Asp4-CSE ↑ |
| MARCKS | Membrane dynamics; cell motility | Asp4-CSE-Asp4 ↑, Asp40-CSE-Asp40 ↑ |
| GSTM3 | Antioxidant defence; prevents oxidative damage | Asp4-CSE-Asp4 ↑ |
| ATPV0D1 | Vacuolar ATPase; endosomal function; EV biogenesis | Asp4-CSE-Asp4 ↑ |

## 4. Discussion

This study characterizes, for the first time, the dose and timing-dependent effects of aspirin on CTCs and their corresponding extracellular vesicle (EV) cargo under oxidative stress conditions that mimic preeclampsia (PE) pathology. CTCs, located at the feto-maternal decidua interface, is known to function as a mechanical barrier, also an immunoregulatory layer, and a major source of pregnancy hormone progesterone. CTC-derived EVs represent key paracrine mediators within the amniochorionic microenvironment, capable of influencing inflammation, angiogenesis, and extracellular matrix (ECM) integrity.^40,28,41^ Using EV proteomics as a molecular readout of cellular status, we demonstrate that low-dose aspirin modulates anti-apoptotic and anti-inflammatory pathways in CTCs in a time and dose dependent manner extending the current understanding of aspirin’s action beyond the placenta, highlighting a secondary but physiologically relevant pharmacological effect on the fetal membrane interface.

### 4.1 CSE Creates a “Pathological EV Signature” of coagulation, apoptosis, and vascular suppression: F10 and MFG-E8 as anchors of a pathological program

The marked upregulation of F10 (Coagulation Factor X) as the single most significantly altered protein in CSE-derived EVs might indicate a direct mechanistic link between CSE exposure and the prothrombotic state characteristic of placental pathology. EV-associated coagulation factors contribute to the thrombo-inflammatory microenvironment of PE^42,43^ and our finding suggests that CTC-EVs from smoking-affected pregnancies actively promote coagulation at the feto-maternal interface. Notably, F10 is detectable by standard clinical coagulation assays, positioning it as an immediately translatable biomarker candidate. The downregulation of MFG-E8 (Lactadherin) may suggest multiple implications. MFG-E8 is known promote efferocytosis, inhibits inflammation, and supports angiogenesis through αvβ3/αvβ5 integrin engagement.^44^ On EVs, MFG-E8 facilitates vesicle target cell interactions. Its depletion from CSE-derived EVs simultaneously impairs anti-inflammatory clearance, pro-angiogenic signaling, and EV uptake efficiency resulting in a compounded loss of protective functions.

The downregulation of VPS4B an ESCRT AAA-ATPase essential for EV biogenesis ^45^ parallels the identification of BROX (also ESCRT-associated) in the ML panel (Figure 11) and suggests CSE disrupts the EV packaging machinery itself. This disruption may explain the coordinated, multi-pathway perturbation observed across the CSE-derived EV proteome The coordinate activation of TNFα/NF-κB, IL-6/JAK/STAT3, p53, interferon-γ, hypoxia, and inflammatory response pathways in CSE-derived EV cargo (Figure 3D) mirrors the molecular landscape of the preeclamptic placenta ^46,47, 48^ suggesting that CTC-EVs from CSE-exposed cells carry a molecular program that, if transferred to maternal cells, could contribute to the systemic inflammatory and endothelial activation characteristic of PE.

### 4.2. Aspirin demonstrates a compartment-specific molecular response within the feto-maternal interface

While aspirin’s vascular and anti-inflammatory benefits in the placenta are well documented, its impact on a critical component of the fetal membrane complex CTC layer, a critical yet distinct component of the fetal membrane complex, has remained uncharacterized. Our proteomics analysis reveals a fundamental functional compartmentalization at the feto-maternal interface. Previous studies have characterized the placenta as the primary pharmacological target, responding to aspirin with angiogenic reprogramming. In contrast, we identify the CTC layer as a distinct’Secondary Responder.’ Unlike placental trophoblasts, which drive vascular remodeling, CTCs respond to prophylactic aspirin by reinforcing structural and survival checkpoints. This suggests that aspirin’s clinical efficacy in preventing preeclampsia is not solely derived from placental perfusion, but also from the stabilization of the chorio-decidual barrier, potentially mitigating the’second hit’ of inflammatory cascade that precipitates preterm birth. This compartmentalized response aligns with the physiological roles of these tissues. The placenta mediates nutrient and gas exchange, demanding distinct angiogenic control, while the chorion primarily maintains intrauterine structural integrity, endocrine functions and immune tolerance. Aspirin’s modest but measurable effects in CTCs may thus serve to maintain immune equilibrium and prevent premature inflammatory activation, a plausible mechanism contributing to its observed reduction of preterm birth risk in clinical settings.

### 4.3 Dose and timing dependent dissociation: A pharmacological window for angiogenic rescue

The finding that only low-dose aspirin restores angiogenesis, while both doses prevent apoptosis, represents the most clinically significant observation of this study, revealing a clear dose-dependent dissociation in CTC-EV responses. Low-dose aspirin selectively inhibits COX-1–mediated thromboxane A_₂_ while sparing COX-2–dependent prostacyclin thereby preserving_₂_-driven trophoblast invasion, vascular remodeling, and angiogenesis, which may sustain pro-angiogenic signaling (LGR4, THY1, CAV1, MMP14, MYOF). In contrast High-dose aspirin inhibits both COX-1 and COX-2, suppressing PGI_2_ as well as TxA ^51^ loss of PGI may negate pro-angiogenic benefits despite anti-inflammatory effects, while residual anti-apoptotic activity likely reflects COX-independent mechanisms (e.g., NF-κB inhibition, antioxidant actions). Low-dose prophylactic aspirin restored a coordinated angiogenic program comprising LGR4 (Wnt/β-catenin activation), THY1/CD90 (adhesion-mediated angiogenesis), CAV1 (VEGFR scaffolding), MMP14/MT1-MMP (ECM remodeling for invasion), and MYOF (VEGFR2 trafficking/signaling), supporting endothelial growth. ^52,53,54,55,56,57,58,59^ When aspirin was administered prophylactically (Asp4-CSE-Asp4) before oxidative stress, CTC-derived EVs showed early suppression of apoptotic mediators (CASP3, BAX) and restoration of ECM-related and angiogenic proteins (MMP14, COL4A1). This pattern suggests that prophylactic aspirin preconditions the chorion, establishing a low-inflammatory and pro cell survival baseline that resists oxidative insult. In contrast therapeutic aspirin (CSE-Asp4) administered post-injury, acted primarily as a rescue mechanism, reducing TNF and p53 activation and stabilizing EV-mediated stress signaling. The ability of post-injury aspirin to restore molecular homeostasis indicates that the drug retains reparative potential, even when prophylactic window has been missed. However, the effects were more limited in scope, consistent with clinical data showing reduced efficacy when aspirin is initiated late in gestation (>16 weeks). High-dose aspirin in both paradigms suppressed not only injury but also adaptive responses, including repair and angiogenic programs, a pattern indicative of over-inhibition of eicosanoid and NF-κB pathways, reinforcing the concept of a narrow therapeutic window.

### 4.4 The Inflammatory divide: prophylaxis modulates, therapy cannot reverse

Perhaps the most important clinical finding is the complete inability of therapeutic aspirin to reverse CSE-established inflammatory programming (Figure 9). Both therapeutic doses showed “No Response Observed” for inflammatory networks, while the heatmap confirmed persistent inflammatory gene activation indistinguishable from untreated CSE. This finding illuminates why aspirin initiated after 16 weeks of gestation has diminished efficacy in clinical trials.^7,9^ By mid-pregnancy, CSE exposure has already established inflammatory programming in trophoblasts. Aspirin initiated at this point can modulate apoptosis and necrosis through COX-independent mechanisms but cannot disassemble the entrenched inflammatory transcriptional state it is, in effect, too late for the inflammatory component of protection. Conversely, prophylactic low dose aspirin particularly in the continuous schedule (Asp4-CSE-Asp4) modulates inflammatory networks (Figures 6, 7) and normalizes the composite Inflammation Index (Figure 9F). This observation suggests that aspirin must be present at the time of or before the inflammatory insult to prevent establishment of the inflammatory program, rather than attempting to reverse it afterward.

Notably, prophylactic co-treatment (Asp+CSE) paradoxically increased the Inflammation Index (Figure 9E-F) whereas continuous treatment normalized it toward baseline. One interpretation is that acute aspirin co-administration during CSE exposure transiently augments inflammatory signaling, but sustained exposure allows the anti-inflammatory program to establish and predominate. This carries direct clinical implications: consistent, continuous aspirin exposure may be more important than the specific timing of initiation carrying direct clinical implications: consistent, continuous aspirin exposure may be more important than the specific timing of initiation.^13,60^

### 4.5 The Therapeutic arm: partial rescue through subtle redistribution

While therapeutic aspirin cannot reverse established inflammatory programming, it does attenuate apoptosis and necrosis (Figure 8) through distributed, modest protein changes rather than the strong reprogramming observed with prophylactic regimen. The therapeutic networks involve many of the same nodes as prophylactic networks (CAPNS1, YBX1, DDX21, FUBP1, MCAM, RAD23B) but with smaller effect sizes. The Supplementary Figure 3 analysis reveals that therapeutic low-dose aspirin (CSE+Asp4) achieves partial rescue but with persistent predicted DNA damage and inflammatory outcomes, while therapeutic high-dose aspirin (CSE+Asp40) produces the most complex network with evidence of both protective and adverse (DNA damage, inflammation of organ) outcomes. Collectively, these findings support a clinical model where therapeutic aspirin offers incomplete protection: sufficient to modulate selected pathological pathways but inadequate for comprehensive rescue, particularly of inflammatory and angiogenic domains.

### 4.6 Toward a Unified model: Timing, Dose, and Duration

Integrating all findings, we propose a three-tiered unified model of aspirin’s effects on CTC-EV cargo, stratified by pharmacological accessibility. Apoptosis prevention: the most broadly accessible effect was achieved by all aspirin regimens at both doses regardless of timing, likely reflecting COX-independent mechanisms including NF-κB modulation, direct protein acetylation, and antioxidant activity. Angiogenic restoration was selectively achieved only by low-dose aspirin, particularly when administered before or during CSE exposure, consistent with preservation of COX-2–dependent prostacyclin signaling within the narrow pharmacological selectivity window afforded by low-dose aspirin. Inflammatory modulation was the most restricted effect, achievable only prophylactically at low dose and requiring continuous exposure for full normalization of the Inflammation Index; once inflammatory programming was established, this effect was completely therapeutically inaccessible. This hierarchical model in which anti-apoptotic protection is universally accessible, angiogenic rescue is dose-restricted, and inflammatory modulation is both dose and timing-restricted provides a mechanistic rationale for why early, continuous, low-dose aspirin affords the most comprehensive protection consistent with the ASPRE trial^7^ and ISSHP guidelines.^64^

### 4.7 Clinical and translational relevance

Clinically, these findings underscore that aspirin’s therapeutic benefit may extend beyond its classical anti-platelet or placental actions. Even subtle modulation of fetal membrane signaling could contribute to improved pregnancy outcomes, particularly in reducing inflammation driven complications such as preterm premature rupture of membranes (PPROM) and spontaneous preterm birth. The dose and timing dependent relationships characterized here, provides a mechanistic rationale for initiating low dose aspirin early in pregnancy to maximize protective efficacy while preserving regenerative and angiogenic processes. Furthermore, the identification of EV-based biomarkers that distinguish aspirin responsive from aspirin non-responsive states opens new avenues for personalized obstetric treatment approaches. Monitoring EV proteomic profiles longitudinally during gestation might help to identify non-responders early guiding timely dosage adjustments or adjunctive interventions.

## 5. Conclusions

This study presents the first comprehensive characterization of CSE effects on CTC-derived EV cargo and the first systematic comparison of prophylactic, therapeutic, and combined aspirin regimens across apoptotic, angiogenic, and inflammatory pathway domains. These EV markers may offer a tangible path toward precision obstetrics, moving beyond the “one-size-fits-all” approach to identify aspirin-responsive pregnancies via non-invasive liquid biopsy. Ultimately, incorporating the chorion into our pharmacological models redefines our understanding of preeclampsia prophylaxis, shifting the focus from simple placental perfusion to holistic feto-maternal barrier defense. Future translational studies integrating PTC and CTC-EV networks, EV profiling, microphysiological models, and maternal plasma biomarker validation will further determine how fetal membrane dynamics affect the overall therapeutic landscape of aspirin in pregnancy.

