## Supplemental figure legend for "From Stress to Survival: Trophoblast-Derived Extracellular Vesicle Proteome Captures Aspirin-Driven Cellular Reprogramming in a Preeclampsia Model"

**Supplementary Figure 1: IPA identifies distinct signaling networks driving prophylactic Aspirin-induced Cell Survival.**

A) IPA molecular interaction network of differentially expressed molecules in CSE vs Ctrl (red, upregulated; green, downregulated).(B) Predicted functional consequences of (A), indicating increased DNA damage, necrosis, apoptosis, and inflammation.(C) IPA molecular network for Asp4 + CSE vs CSE, showing treatment-induced modulation of affected molecules.(D) Predicted functional outcomes of (C), demonstrating reduced or absence of necrosis, apoptosis, and inflammatory pathways following Asp4 treatment.(E) Integrated molecular network of Asp40 + CSE vs CSE, highlighting combined effects of exposure and treatment on regulatory pathways.(F) Predicted functional consequences of (E), summarizing overall changes in injury, reduced necrosis, apoptosis, and inflammatory responses relative to control.

Key: Red and green nodes represent upregulated and downregulated proteins, respectively. Orange lines indicate predicted activation; blue lines indicate predicted inhibition of downstream targets. Solid lines represent direct interactions; dashed lines represent indirect interactions.

**Supplementary Figure 2. Sequential aspirin treatment (pre- and post-CSE) reprograms DNA damage, cell death, and inflammatory signaling.** (A) IPA molecular interaction network: Asp40-CSE vs. CSE. Features include RAS-related signaling, tyrosine kinase networks, MARCK3, VAPA/VAPB, TNF regulation, and CRADD/SLC2A8 nodes. (B) Predicted functional consequences of (A): absence of DNA damage, reduced apoptosis and inflammatory signaling, but increased necrosis — a potential adverse effect of high-dose aspirin co-treatment. (C) IPA molecular network: Asp40-CSE-Asp40 vs. CSE. Features include GPCR-ligand complexes, GANT/GANKB complexes, PTPRM/PTPRS interactions, and PIK4KA signaling. (D) Predicted functional outcomes of (C): absence of DNA damage and inflammation-related processes, reduction in apoptosis. The distinct network architectures between Asp40-CSE and Asp40-CSE-Asp40 conditions indicate that extended high-dose aspirin exposure activates different molecular pathways than co-treatment alone.

**Supplementary Figure 3. Therapeutic aspirin mitigates CSE-induced cellular injury via network-level reprogramming.** (A) IPA molecular interaction network: CSE+Asp4 vs. CSE. Features include TP53 (p53), EP300, DSCR2, EPB42, TMEM106B, MCAM, STX17, TFEB, VAMP8, YTHDF1, GALNT3, and endosomal/autophagy components. (B) Predicted functional consequences of (A): DNA damage and inflammatory outcomes persist alongside reduced apoptosis, indicating incomplete rescue. (C) IPA molecular network: CSE+Asp40 vs. CSE. The most complex network in the study, features MAP3K4, BLNK, SNX1, CDK7, TP73, RPS6KA3, ARHGEF12, ABCC1, PLD1, CEBAB, RALY, ARCE1, and extensive multi-pathway engagement. (D) Predicted functional consequences of (C) reduced necrosis and apoptosis, but increased DNA damage and inflammation-related pathways (inflammation of organ predicted). High-dose therapeutic aspirin therefore exhibits a dual character protective against cell death but insufficient to suppress injury-associated programs. Node color: red/pink = upregulated; green = downregulated; blue = predicted inhibition. Edge color: orange = activation; blue = inhibition; yellow = inconsistent with state.
