## Supplementary figures and images for "From Stress to Survival: Trophoblast-Derived Extracellular Vesicle Proteome Captures Aspirin-Driven Cellular Reprogramming in a Preeclampsia Model"

### Supplementary Figure 1

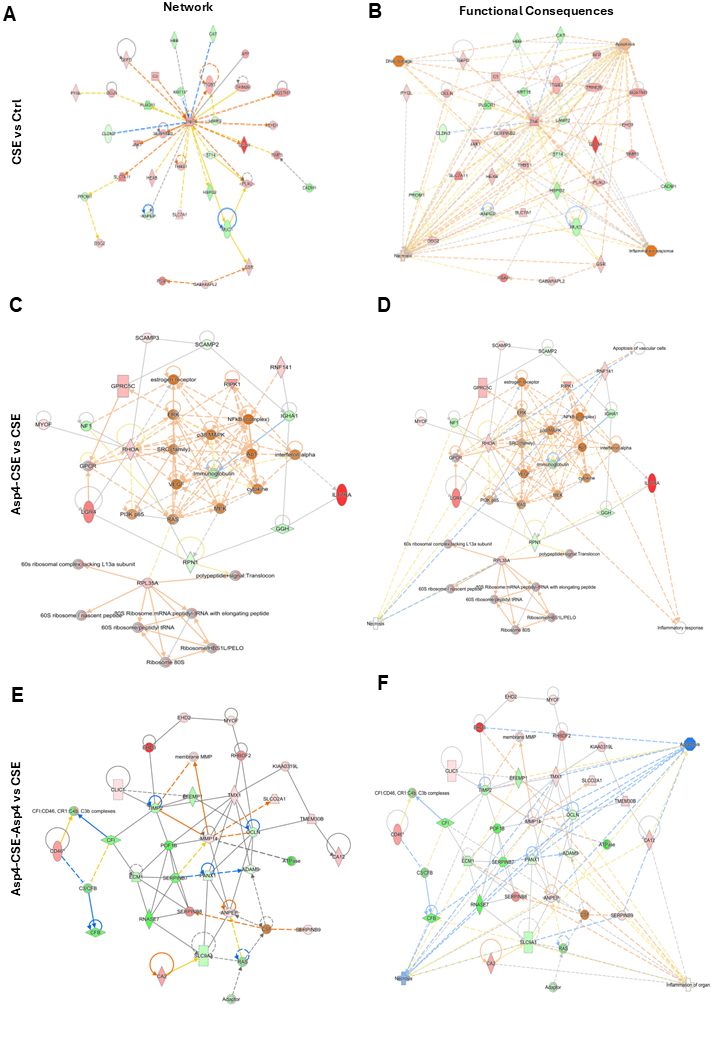

### Supplementary Figure 2

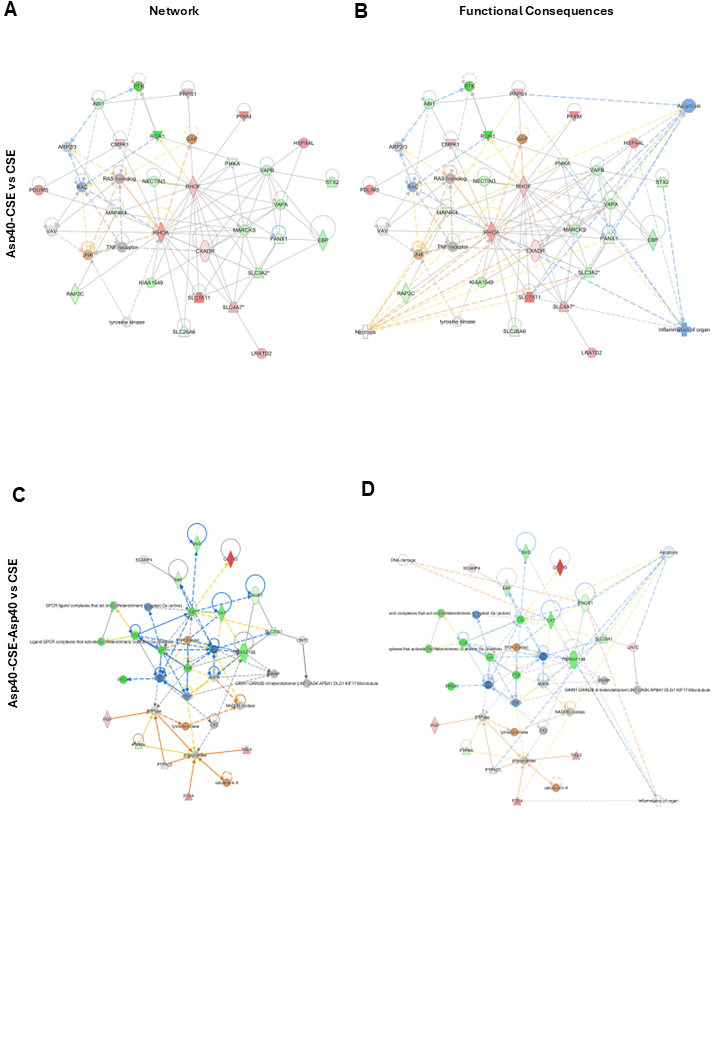

### Supplementary figure 3

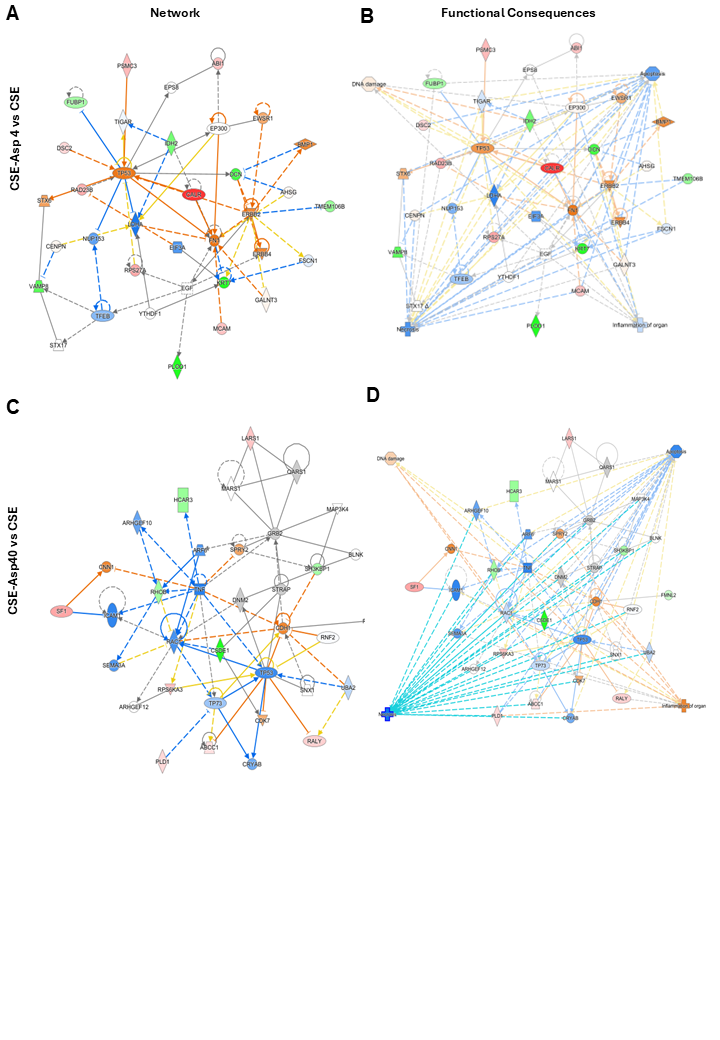
